# Heterogeneous lineage-specific arginine deiminase expression within dental microbiome species

**DOI:** 10.1101/2023.12.07.570591

**Authors:** Allison E. Mann, Brinta Chakraborty, Lauren M. O’Connell, Marcelle M. Nascimento, Robert A. Burne, Vincent P. Richards

**Affiliations:** Department of Biological Sciences, Clemson University, Clemson, SC, USA; Department of Oral Biology, College of Dentistry, University of Florida, Gainesville, FL, USA; Department of Restorative Dental Sciences, Division of Operative Dentistry, College of Dentistry, University of Florida, Gainesville, FL, USA

**Keywords:** metatranscriptomics, arginine deiminase system, probiotics, oral microbiome, caries

## Abstract

Arginine catabolism by the bacterial arginine deiminase system (ADS) has anticariogenic properties through the production of ammonia, which modulates the pH of the oral environment. Given the potential protective capacity of the ADS pathway, the exploitation of ADS competent oral microbes through pre- or probiotic applications is a promising therapeutic target to prevent tooth decay. To date, most investigations of the ADS in the oral cavity and its relation to caries have focused on indirect measures of activity, or on specific bacterial groups, yet the pervasiveness and rate of expression of the ADS operon in diverse mixed microbial communities in oral health and disease remains an open question. Here we use a multivariate approach, combining ultra-deep metatranscriptomic sequencing with paired metataxonomic and *in vitro* citrulline quantification to characterize the microbial community and ADS operon expression in healthy and late-stage cavitated teeth. While ADS activity is higher in healthy teeth, we identify multiple bacterial lineages with upregulated ADS activity on cavitated teeth that are distinct from those found on healthy teeth using both reference-based mapping and *denovo* assembly methods. Our dual metataxonomic and metatranscriptomic approach demonstrates the importance of species abundance for gene expression data interpretation and that patterns of differential expression can be skewed by low abundance groups. Finally, we identify several potential candidate probiotic bacterial lineages within species that may be useful therapeutic targets for the prevention of tooth decay and propose that the development of a strain-specific, mixed-microbial probiotic may be a beneficial approach given the heterogeneity of taxa identified here across health groups.

**IMPORTANCE:** Tooth decay is the most common preventable chronic disease, globally affecting more than two billion people. The development of caries on teeth is primarily a consequence of acid production by cariogenic bacteria that inhabit the plaque microbiome. Other bacterial strains in the oral cavity may suppress or prevent tooth decay by producing ammonia as a byproduct of the arginine deiminase metabolic pathway, increasing the pH of the plaque biofilm. While the benefits of arginine metabolism on oral health have been extensively documented in specific bacterial groups, the prevalence and consistency of ADS activity among oral bacteria in a community context remains an open question. In the current study, we use a multi-omics approach to document the pervasiveness of expression of the ADS operon in both health and disease to better understand the conditions in which ADS activity may prevent tooth decay.

## INTRODUCTION

Arginine catabolism is a common metabolic pathway for anaerobic energy production used by a variety of organisms including bacteria, archaea, and microbial eukaryotes (Novák et al. 2016a). Through a series of enzymatic modifications, arginine ingested from food, endogenously produced by the host, or synthesized by resident bacteria, is broken down by the arginine deiminase system (ADS) to produce citrulline, ornithine, carbon dioxide, ammonia, and ATP (Burne and Marquis 2000; Cunin et al. 1986; Nascimento, Alvarez, Huang, Hanway, et al. 2019). Along with acting as an important source of energy, carbon, and nitrogen production, the ADS pathway is protective against low pH environments through the production of ammonia (Burne and Marquis 2000; Marquis et al. 1987; Nascimento et al. 2014). Furthermore, research has demonstrated that ADS activity affects the assembly and coaggregation of bacteria in the plaque biofilm matrix, prevents the colonization of the cariogenic bacterium *Streptococcus mutans*, and raises the pH of the oral environment, all of which may inhibit caries development (Chakraborty and Burne 2017; He et al. 2016; Kamaguchi et al. 2001; Nascimento et al. 2009; Nascimento et al. 2013; Sharma et al. 2014). Moreover, growth of bacteria in the presence of arginine *in vitro* or administration of arginine orally *in vivo* increases both ADS activity and the pH of the oral environment, and numerous clinical trials have documented a decrease in caries incidence when an arginine-supplemented toothpaste was used (Acevedo et al. 2005, 2008; Carda-Diéguez, Moazzez, and Mira 2022; Nascimento et al. 2014; Wolff and Schenkel 2018). Given the capacity of ADS activity to buffer the effects of highly acidogenic microbes, ADS competent bacteria that inhabit oral biofilms (i.e., dental plaque) have been extensively studied in the context of oral health and the prevention of tooth decay (Casiano-Colón and Marquis 1988; Nascimento et al. 2014; Nascimento and Burne 2014; Nascimento 2018; Reyes et al. 2014). Of particular interest are the mechanisms of ADS pathway regulation in streptococci, many of which are important early colonizers of the oral cavity. (Curran et al. 1998; Dong et al. 2002a; Ferro, Bender, and Marquis 1983; Griswold et al. 2004; Poolman, Driessen, and Konings 1987; Quivey, Kuhnert, and Hahn 2001). Arginine and ADS competent oral symbionts may therefore be candidate pre- and probiotic treatments, respectively, yet little is known about ADS activity among members of the oral cavity in a mixed microbial community context.

In the current study, we used a multivariate metatranscriptomic, metataxonomic, and *in vitro* assay approach to characterize the composition and function of the supragingival plaque microbial community in health and late-stage caries development. We found that while some *Streptococcus* sp. have higher expression of the ADS operon in health (i.e., *S. oralis* and *S. sanguinis*), others are more highly expressed in disease (e.g., *S. parasanguinis*). Importantly, upregulation of genes involved in ADS activity for diseased teeth communities is limited to those with a low abundance of *Streptococcus mutans*, which may indicate an antagonistic relationship between these groups. Moreover, we find diverse non-*Streptococcus* bacteria with higher ADS operon expression in health when compared to disease. For instance, two poorly characterized members of the genus *Leptotrichia* have relatively high ADS expression in health and may represent an untapped source of genetically diverse potential probiotic strains. In addition, our results highlight the importance of community composition and within-species, lineage, or strain-level resolution in the interpretation of metatranscriptomic data from a mixed microbial community. We found that community composition is significantly correlated with observed differences in ADS gene expression profiles and that, within species, only some lineages: (1) possess a full ADS operon, (2) have relatively high community abundance, and (3) have high ADS expression, all of which have important implications for the development of a probiotic. Moreover, comparing our reference-database mapping approach to that generated using a *denovo* assembly approach documents substantial genetic diversity within lineages that is not well represented in publicly available databases (e.g., HOMD). Finally, we calculated coverage across the ADS operon in a subset of species and found that post-transcriptional regulatory activity may affect the efficacy of some strains to modulate the pH of the oral environment, a pattern which is substantiated by *denovo* assembly of full-length ADS transcripts in our dataset. Assessment of the relative contributions of transcriptional versus post-transcriptional control would require extensive strain-level analyses. Finally, while the aim of this study is to characterize the microbial community and ADS activity in health and disease, the vast majority of observed functional variation between healthy teeth and those with carious lesions is driven by diverse and discordant gene expression among communities associated with severe tooth decay. The results of this study highlight the functional and taxonomic volatility in late-stage caries development, identifies alternate microbial candidates for probiotic development in the context of the ADS pathway, and highlights the importance of accounting for community composition when interpreting differential gene expression in mixed microbial contexts.

## RESULTS

### Increased microbiome diversity and citrulline production in health

We collected 60 supragingival plaque samples from 54 adults using a standard protocol (see Materials and Methods). Multiple teeth were sampled from a subset of individuals (n=3) to document intraindividual functional variation. Each plaque sample was classified as either originating from a caries-free tooth (PF) or a tooth with an active, dentin cavitation or caries lesion (PD). In total, 27 plaque samples were collected from teeth with an active dentin cavity (PD) and 33 from a caries-free tooth (PF). Study participants included 35 females and 25 males ranging in age from 18 to 65 years old (average 31 ± 11 years).

First, to characterize the diversity of the oral microbiome, and account for differences in community composition driving gene expression, we used a high-resolution metataxonomic approach targeting a fragment of the bacterial *rpo*C gene to profile the community composition of each sample. We chose this approach as (1) it is more cost-effective than metagenomic sequencing, (2) the *rpo*C gene is a single-copy core gene in bacteria and therefore should be a better representation of community membership as compared to other commonly used marker genes (e.g., 16S rRNA hypervariable regions), and finally, (3) the *rpo*C gene has high phylogenetic resolution at the species and strain level (Hassler et al. 2022) (see Materials and Methods), which allows us to directly compare community composition to gene abundance data using a reference-based mapping approach. These data were compared to citrulline assay and metatranscriptomic results generated from the same samples to clarify the role of community composition and membership in ADS gene expression in health and disease using both biochemical and RNA sequencing techniques.

We found that the oral microbiome of healthy teeth (PF) is more diverse than those with caries (PD) as measured by Shannon diversity (Wilcoxon rank sum test: p=0.003; Fig. 1a), and we found that citrulline production is significantly higher in PF as compared to PD samples (Wilcoxon rank sum test: p=0.0004). Variation in citrulline production is significantly (albeit weakly) correlated with beta diversity when controlling for tooth type (PD vs. PF) (PERMANOVA: R^2^=0.03; p=0.03) (Fig. 1b). Importantly, however, not all PF samples have high citrulline production nor do all PD samples have low production. For example, a single PD sample (UF68PD) had unexpectedly high citrulline production (Fig. 1b) which may be the consequence of significantly higher ADS activity by *Streptococcus oralis* as detected by metatranscriptomic sequencing (see below) (Fig. 1c, Table S1). Specifically, although the proportion of *S. oralis* assigned to this sample is small in our paired metataxonomic dataset (<1% of the total community), transcripts from *S. oralis* make up 29.56% of all ADS transcripts originating from UF68PD. The median proportion of transcripts assigned to this species in PF is 12.44% (± 13.17%) while in PD it is 1.80% (± 15.91%) making UF68PD more similar to other high citrulline producing PF samples in terms of the proportion of transcripts assigned to this species. This is in stark contrast to four low-citrulline producing PF samples (UF33PF, UF36PF, UF40PFR, UF48PF) where the proportion of transcripts assigned to *S. oralis* was less than 1% each. In addition to *S. oralis*, sample UF68PD also had a high number of ADS gene transcripts assigned to *Leptotrichia* sp. oral taxon (HMT) 215, *Kingella oralis*, and *Actinomyces naeslundii* (Table S2) which were also found to have higher ADS expression in PF as compared to PD.

**FIG 1.**
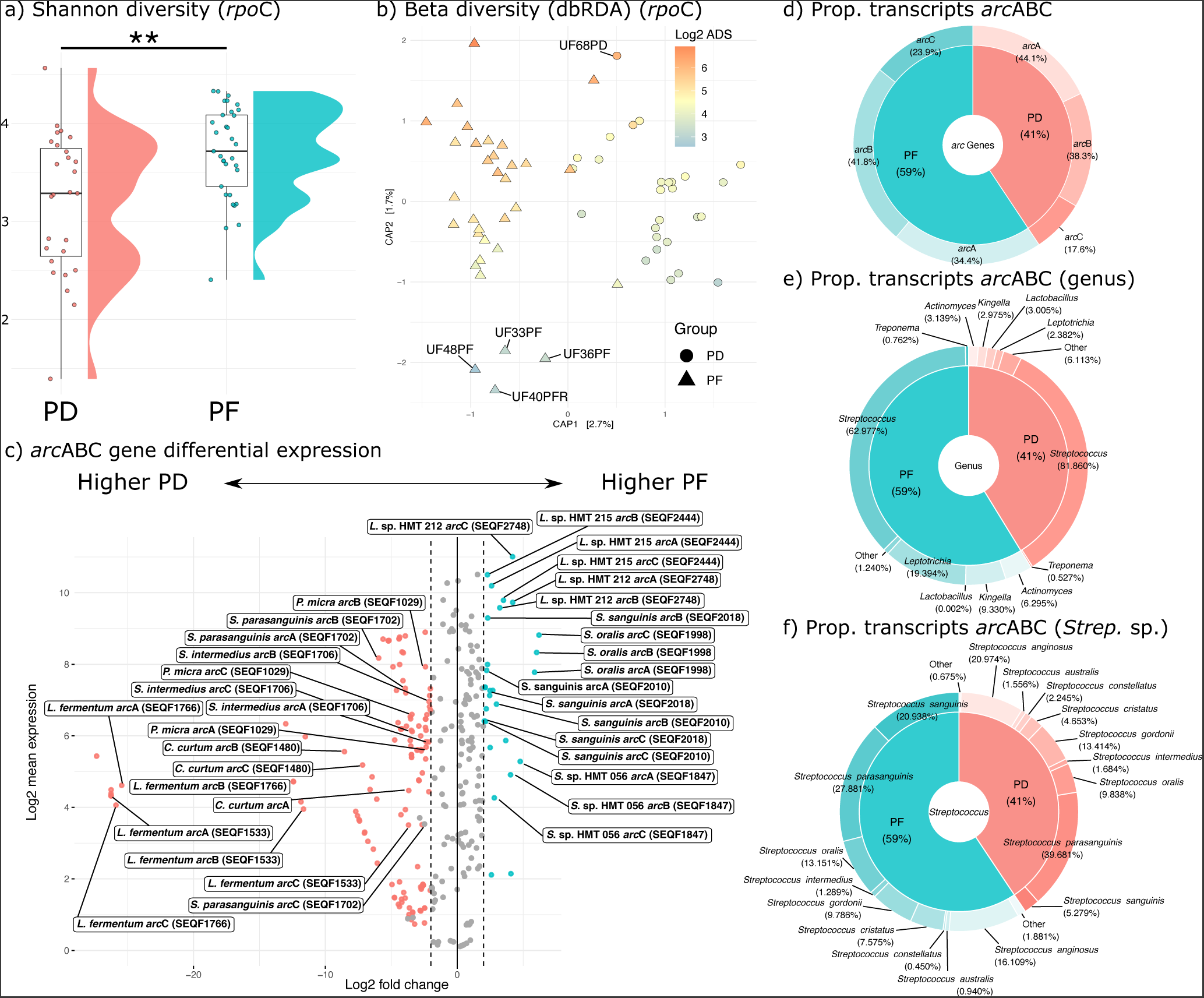
Significant differences in ADS activity found comparing PD and PF are predominantly driven by major differences in taxonomic composition: (a) Shannon diversity box and raincloud plots for PD and PF samples. (b) Beta diversity colored by ADS expression levels as measured by our citrulline assay. Shapes represent the two groups. Samples with unexpectedly high or low citrulline production labelled on plot (see text). (c) Differential abundance volcano plot of metatranscriptome data. Colored points represent genes that were both statistically significant and passed log fold change abundance thresholds (see text). Labeled points represent all taxa upregulated for PF and six select taxa upregulated for PD (see text) that (i) had the highest adjusted p-value for either PD (pink) or PF (blue) and (ii) had all three *arc* genes marked as differentially abundant. One exception shown here is *S. sanguinis* genes *arc*C1 and *arc*B which, despite having significantly higher expression of the *arc*A gene, did not pass thresholds. HOMD strain identifiers along with species identifications are shown in boxes. (d) Proportion of reads mapping to each of the three major *arc* genes in the ADS operon after manual curation for genes that were present in synteny within a contig in both PF and PD samples. (e) The proportion of RNA seq reads that mapped to *arc*ABC genes among different genera. (c) Proportion of reads that mapped to *arc*ABC across different *Streptococcus* species (strains collapsed to species).

### Unequal expression of individual *arc*ABC genes

While arginine catabolism is a common metabolic pathway found across diverse bacterial groups, it is dispensable and the operon is absent or truncated in many oral bacteria, even among closely related phylogenetic groups (Table S3) (Tian et al. 2022). Moreover, gene content and organization of the ADS operon varies within and between ADS competent bacteria (Dong et al. 2002b; Noens and Lolkema 2016a; Velsko et al. 2018; Zúñiga et al. 1998; Zúñiga, Miralles, and Pérez-Martínez 2002). As such, we first identified ADS competent oral taxa in the Human Oral Microbiome Database (HOMD) (Chen et al. 2010) through homologous gene clustering of three highly conserved *arc* genes that produce enzymes necessary for ADS activity (1) *arc*A (arginine deiminase EC: 3.5.3.6), (2) *arc*B (ornithine carbamoyltransferase EC: 2.1.3.3), and (3) *arc*C (carbamate kinase EC: 2.7.2.2). Only those bacteria with all three genes in synteny were included in differential gene expression analyses in our reference based mapping approach (see Materials and Methods). Using this manually curated collection of ADS operons as a reference, we next compared the relative abundance of transcripts assigned to individual *arc* genes across PF and PD samples. As only those bacteria with *arc*ABC organized in a syntenous operon were considered, we expected the proportion of transcripts that mapped to each gene within an operon to be approximately equivalent to one another. Interestingly, for both PF and PD samples, while the proportion of transcripts assigned to *arc*A and *arc*B were similar, the proportion of transcripts mapping to *arc*C tended to be lower. This is especially true of transcripts mapping to the *arc* genes in PD samples where the proportion for *arc*A is nearly three times higher than *arc*C (Fig. 1d). This may be due to methodological biases (e.g., incomplete or scaffold level genome assemblies used as reference), but also may be indicative of differences in operon usage or structure in ADS competent bacteria that are potentially associated with health status. For example, specific genes within an operon may be regulated through mRNA degradation depending on the transcriptional needs of the cell (Belasco et al. 1985; Dar and Sorek 2018; Newbury, Smith, and Higgins 1987), internal transcription promoter or termination sites, or through other forms of gene expression regulation (e.g., transcriptional attenuation through secondary DNA structures that prevent transcription of the full operon) (See Güell et al. 2011). Notably, for this analysis, we only considered reference operons that were (1) complete (i.e., all three genes present) and (2) present on the same contig (to reduce the impact of incomplete genomes) which suggests these patterns are biological and not methodological in nature.

Of those transcripts that mapped to one of the three *arc* genes analyzed here, most were assigned to *Streptococcus,* but we also detected ADS *arc* gene expression for other genera including *Leptotrichia, Kingella, Lactobacillus, Actinomyces, Treponema* and several low frequency groups (Table S2; Fig. 1e). While both PD and PF samples had a high frequency of transcripts assigned to *Streptococcus parasanguinis* (PF: 27.88%, PD: 39.68%) and *S. anginosus* (PF: 16.11%, PD: 20.97%), PF samples had a comparatively higher proportion of transcripts assigned to *S. sanguinis* (PF: 20.94%, PD: 5.28%) and *S. oralis* (PF: 13.15%, PD: 9.84%) as compared to PD (Fig. 1f). To better understand how structure and regulation might impact the production and survival of transcripts, we mapped our quality filtered metatranscriptomic data to the full ADS operon of six strains of interest (*Streptococcus sanguinis* SEQF2018, *S. parasanguinis* SEQF1919, *S. gordonii* SEQF1066, *S. oralis* SEQF1998, *Leptotrichia* oral taxon 212, and *Leptotrichia* oral taxon 215) and generated coverage plots (Fig. 2). We chose these specific taxa as they demonstrate the range of coverage patterns and intergenic region size variation we observed in the dataset across different ADS competent species. Operon coverage for *S. oralis* and both lineages of *Leptotrichia* exhibits a wave-like pattern, with inconsistent coverage levels across the full operon, while coverage for *S. sanguinis*, *S. parasanguinis*, and *S. gordonii* is wave-like in some regions but relatively consistent in others, particularly across *arc*B and to a lesser extent, *arc*A. Interestingly, we found that some of the most precipitous drops in coverage correspond to intergenic regions in the operon, the structure of which varies across the selected taxa presented here. For example, while *S. sanguinis, S. parasanguinis*, *S. oralis,* and *S. gordonii* have an intergenic region between *arc*A and *arc*B spanning approximately 51 to 85 base pairs long, both *Leptotrichia* lineages have an intergenic region between *arc*A and *arc*B spanning approximately 142 to 146 base pairs. Additionally, all *Streptococcus* sp. included in this analysis have an intergenic region ranging from between 86 base pairs (*S. parasanguinis*) to 313 base pairs (*S. oralis*) separating *arc*B and *arc*C. In contrast, both *Leptotrichia* sp. have only two or three base pairs separating the two genes. Importantly, the two most commonly observed species, *S. sanguinis* and *S. parasanguinis,* have markedly lower *arc*C coverage as compared to *arc*A and *arc*B which may indicate differential regulation of this gene post transcription. It is possible that drops in coverage for these two species correspond to hotspots or promoters in the operon for transcript degradation and intergenic regions may play a functional role in post-transcriptional regulation of the operon. Notably, the heterogeneity in operon structure and mRNA abundance observed in this study is consistent with the knowledge that ADS expression is known to be complex and responsive to a variety of environmental inputs (e.g., Liu et al. 2008a). A full table of *arc* operon coordinates and lengths of *Streptococcus* sp. found in Fig. 2 and the two *Leptotrichia* sp. analyzed here can be found in Table S4.

**FIG 2.**
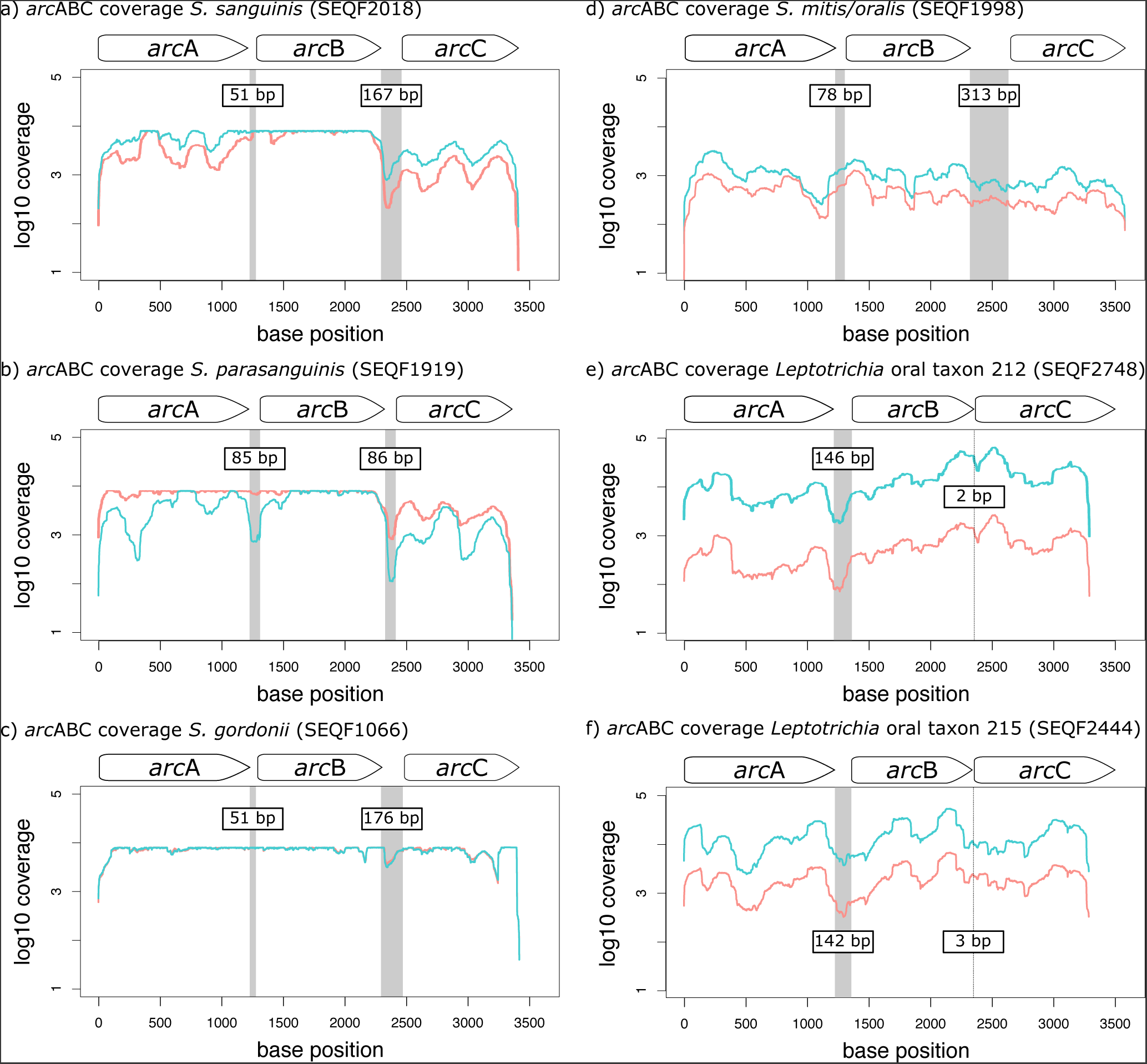
ADS operon structure and coverage varies across strains. (a) Log 10 coverage per base of the three *arc* genes in *Streptococcus sanguinis* (SEQF2018). (b) Log 10 coverage per base of the three *arc* genes in *Streptococcus parasanguinis* (SEQF1919). (c) Log 10 coverage per base of the three *arc* genes in *Streptococcus gordonii* (SEQF1066). (d) Log 10 coverage per base of the three *arc* genes in *Streptococcus mitis/oralis* (SEQF1998). (e) Log 10 coverage per base of the three *arc* genes in *Leptotrichia* sp. oral taxon 212 (SEQF2748). (f) Log 10 coverage per base of the three *arc* genes in *Leptotrichia* sp. oral taxon 215 (SEQF2444). Line colors in all coverage plots represent all reads mapped to health (blue) as compared to disease (red). Grey bars indicate intergenic regions. White internal labels on intergenic regions indicate the size of each region.

### Strain-level differential expression of the ADS operon

Given the heterogeneity of ADS operon structure and presence or absence among closely related oral bacteria, we performed differential expression analyses at both the species level, defined as the sum total of all transcripts that aligned to a named oral species in the HOMD database, and at the strain level, defined as those transcripts that aligned to a particular reference genome within the HOMD database or to a unique reference sequence generated using *denovo* assembly (see below). We take this tiered approach as previous research has demonstrated that ADS activity can vary significantly among closely related oral bacteria (Velsko et al. 2018) which has implications for the development of probiotic treatments. Importantly, however, the HOMD database based strain perspective has limitations. Specifically, a strain within the database used as reference may not be present within a sample. Consequently, when a sequence read aligns to a specific genome within the database, we interpret this as expression (RNA) or presence (DNA) of a strain from the same lineage as the strain in the database and this is understood when we refer to strains throughout the manuscript.

First, we performed differential gene expression analysis to identify strains where all three ADS genes were significantly upregulated in health as compared to disease. Interestingly, only six bacterial strains met these criteria in health including *Streptococcus* sp. oral taxon 056 (SEQF1847), *S. oralis* (SEQF1998), *S. sanguinis* (SEQF2018, SEQF2010), *Leptotrichia* sp. oral taxon 215 (SEQF2444), and *Leptotrichia*. sp. oral taxon 212 (SEQF2748), while 13 were found upregulated in disease including *Parvimonas micra* (SEQF1029), *Cryptobacterium curtum* (SEQF1480), *Limosilactobacillus fermentum* (SEQF1766, SEQF1533, SEQF1964, SEQF2541, SEQF2605), *S. parasanguinis* (SEQF1702, SEQF1706, SEQF1885, SEQF1919, SEQF2007, SEQF2222, SEQF2344, SEQF2625), *S. intermedius, Actinomyces graevenitzii* (SEQF2031, SEQF2450), *Lactobacillus crispatus* (SEQF2084, SEQF2421), *Cutibacterium acnes* (SEQF2166), *S. constellatus* (SEQF2211, SEQF2589, SEQF2590, SEQF2591, SEQF2643), *S. anginosus* (SEQF2334, SEQF2587, SEQF2588, SEQF2609), *Olsenella uli* (SEQF2631), *Olsenella* sp. oral taxon 807 (SEQF2751), and *Lactobacillus ultunensis* (SEQF2852) (Table S1; Fig. 1c). Genes of all six significant strains in PF and a subset of six significant strains in PD (as determined by the highest adjusted FDR score) are highlighted in Fig. 1c.

### Species-level differential expression of the ADS operon

Next, as some species in our reference database have multiple representative strains (predominantly *Streptococcus*), and some strains were more highly expressed than others in the above analyses, we collapsed our results by species designation to compare to strain-level results (Table S5). While most differential expression results remained significant between groups after collapsing at the species level (n = 16), 11 were no longer identified as having higher or lower expression in either group (Table S5 & S6), reflecting strain-level gene expression heterogeneity.

To compare our species and strain-level results further, we next built a maximum likelihood phylogeny using a concatenation of the ADS genes (*arc*A, *arc*B, *arc*C) for all *Streptococcus* species detected in our metatranscriptomic data and identified strains within species that had higher or lower expression or abundance among health groups. For example, while there are a total of 16 representative genomes for *Streptococcus oralis* in HOMD (as of database version 9.15a) and we assigned transcripts to four of them (Fig. 3), only a single *S. oralis* strain had all three genes significantly upregulated in health (SEQF1998) and collapsing at the species level produced an insignificant adjusted (FDR) p value (0.8) (Table S6 & S7). Similarly, while there are eight representative strains in HOMD for *S. intermedius* and all eight had mapped transcripts in our dataset, only one had higher ADS gene expression in PD samples (SEQF1706) and the collapsed data was insignificant (FDR = 0.07). Importantly, unlike other *Streptococcus* groups where average log fold change and base mean are relatively identical (indicating that most transcripts assigned to those lineages mapped equally well to other members of the clade), all *S. oralis* identified in the current data had high average base mean but only SEQF1998 had high average log fold expression (Fig. 3).

**FIG 3.**
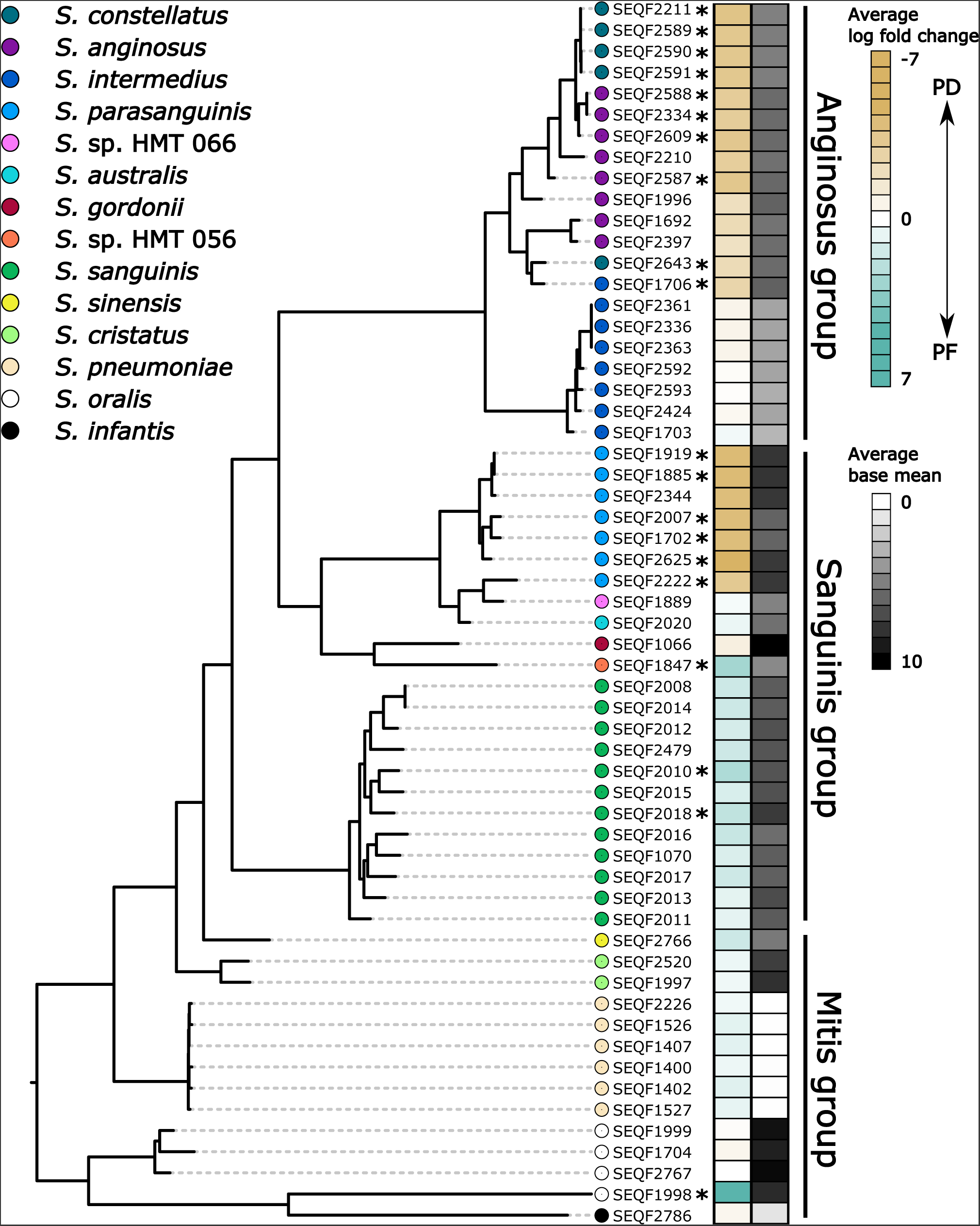
Differential expression and abundance of ADS operon varies by *Streptococcus* lineages. Maximum likelihood phylogeny for the *Streptococcus* sp. ADS operon (concatenated *arc*A, *arc*B, *arc*C genes). Asterix (*) indicates a lineage where all three *arc* genes were up or down regulated as measured by DESeq2 analysis. Corresponding heatmap indicates the average log fold change and base mean per aligned lineage.

### ADS transcript abundance and species abundance are correlated

Because we anticipate community structure to be an important confounding variable when comparing gene expression activity between PF and PD samples (Fig. 1b), we next calculated the correlation between log fold change of the abundance of ADS competent species as measured by *rpo*C amplicon sequencing to the log fold change of gene expression for the ADS operon as measured by our reference based mapping approach (Table S8). By inferring the taxonomic origin of individual transcripts and amplicon sequences with the same reference database, we can then directly compare patterns of community abundance and gene expression levels for specific bacterial species. In this way, we can better understand how community membership and abundance affects our interpretation of gene expression data. Log fold change for ADS expression was calculated at the species-level by first summing all read counts per gene for each HOMD species and then averaged over the three *arc* genes to account for differences in individual gene expression levels. We found that fold change of ADS expression and species abundance was highly correlated (R^2^=0.62, p=0.001), and in most cases, species with high differential expression levels are also found at higher abundance in their respective health category. For example, while transcripts assigned to the ADS pathway in *Streptococcus sanguinis* are more highly abundant among healthy samples, the relative abundance of ASVs assigned to *S. sanguinis* is also higher in health which suggests that the high abundance of transcripts assigned to the ADS pathway is linked to a higher abundance of this species in health as compared to disease. Only 48.15% of PD samples had at least one *rpo*C amplicon assigned to *S. sanguinis* while the species was observed among all PF samples. Moreover, among those samples with detected levels of *S. sanguinis*, it was more highly abundant in PF as compared to PD (PF: 3.7% ± 4.5% relative abundance, PD: 0.5% ± 1.0% relative abundance). Similarly, while transcripts assigned to the ADS pathway in *S. parasanguinis* are more highly abundant in PD, the relative abundance of *S. parasanguinis* is also higher in PD as compared to PF when the amplicon data is considered (average 1.1% ± 2.3% of the total microbial community as compared to 0.4% ± 0.9% of the total community among PF) (Figs. 4a, 4b, 4c). This is despite the fact that there were an approximately equivalent proportion of PF and PD samples with *rpo*C reads assigned to *S. parasanguinis* (52.85% and 60.61%, respectively). Interestingly, we found a subset of ADS competent bacteria that did not follow a simple pattern of higher community abundance coupled with higher ADS transcript expression. For example, *Bulleidia extructa, S. constellatus,* and *S. intermedius* are more highly abundant in PF as compared to PD samples but have higher ADS expression in PD. Conversely, *Cryptobacterium curtum, Parvimonas micra*, and *S. anginosus* have higher ADS expression in PD with very little difference in amplicon abundance between PD and PF samples. While these taxa have high ADS activity in disease that diverges substantially from amplicon sequence variant (ASV) abundances, they are also relatively rare across individuals in this dataset (Fig. 4a) and therefore may be less robust to different oral environments.

**FIG 4.**
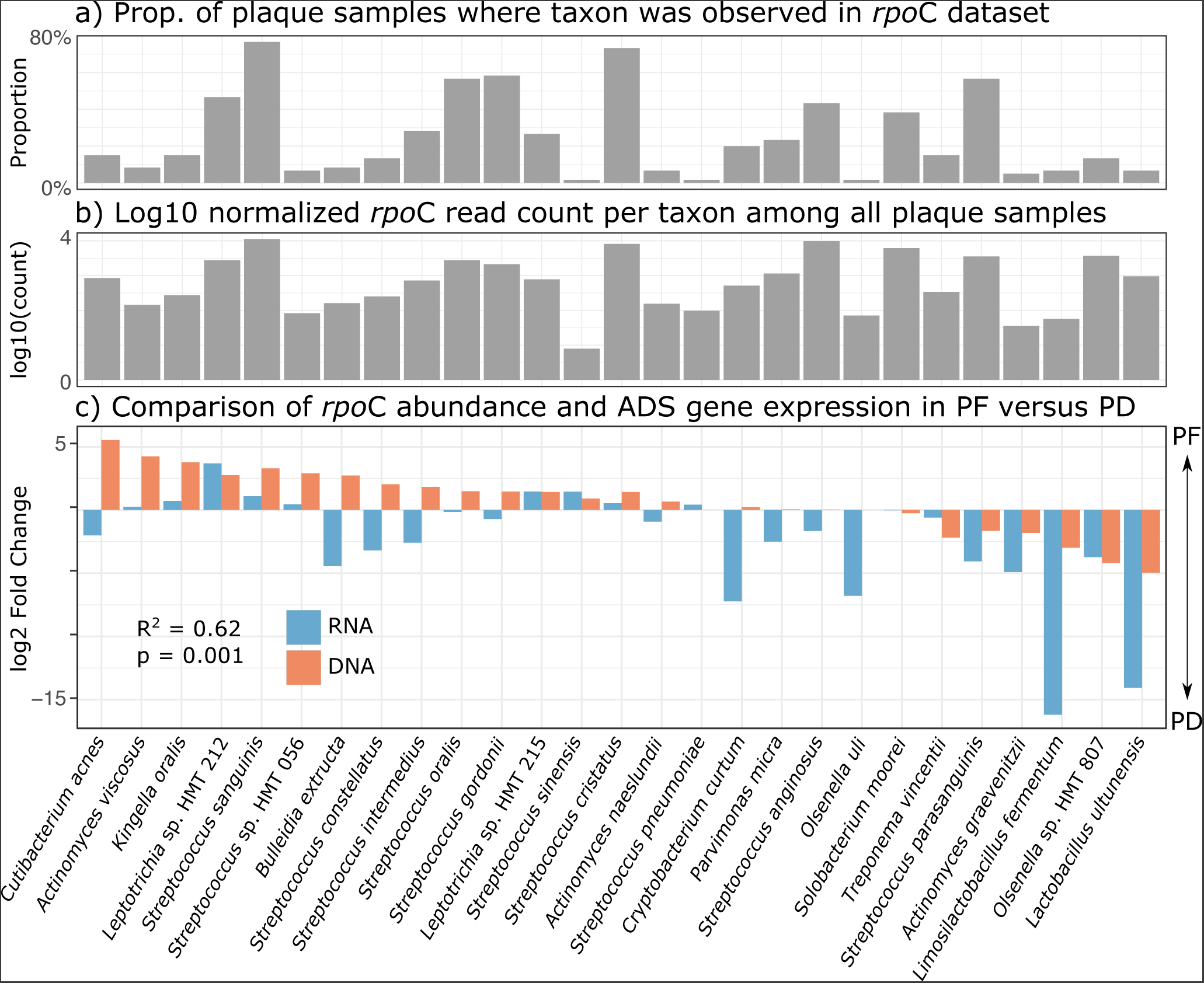
Species level ADS operon expression is significantly correlated to community composition as measured by *rpo*C amplicon sequencing: (a) The proportion of samples in the full amplicon dataset where species listed in panel c were detected. (b) Log 10 normalized read count of amplicon sequences for each species corresponding to panel c. (c) Comparison of log2 fold changes for ADS competent bacteria in the metataxonomic and metatranscriptomic dataset. R^2^ and alpha significance value calculated using a Pearson correlation test.

### *Denovo* assembly of transcripts provides higher ADS strain resolution

As described above, an ongoing challenge in microbial metatranscriptomics is that, lacking other evidence (e.g., metataxonomics or whole genome sequencing), the community of microorganisms across samples is largely unknown. Moreover, the composition of publicly available genomic databases tends to be heavily skewed towards model or pathogenic organisms. For example, of the 555 genomic sequences deposited in HOMD v9.15a for *Streptococcus*, 45% are the cariogenic pathogen *S. mutans*. Conversely, only 16 *Leptotrichia* genomes are available for reference within the database. As such, reference based metatranscriptomic analysis underestimates true strain and species level diversity in gene expression. Given the limitations of a reference-based mapping approach to characterize the full functional profile of a metatranscriptomic dataset of unknown community composition, we next performed an additional *denovo* assembly of our dataset to document the diversity of ADS transcripts across all taxonomic groups.

On average, we generated 364,811 (± 159,936) assembled transcripts per sample (Table S9) of which 1,723 were annotated with at least two of the *arc*ABC genes and 568 had all three genes annotated in synteny. Most transcripts were truncated which supports our previous interpretation of post-transcriptional regulation of the operon using our reference-based mapping approach. Of those ADS transcripts with all three genes present, 132 had one or more genes upregulated in disease (PD) consisting mostly of transcripts assigned to *Streptococcus* sp. (35%) followed by *Lactobacillus ultunensis* (11%) and *Limosilactobacillus fermentum* (8%). Comparatively, 252 unique transcripts were upregulated in health (PF), 32% of which were assigned to *Leptotrichia* sp. oral taxon 212 followed by *Streptococcus* sp. (27%) and *Kingella oralis* (18%) (Table S10). The results of this analysis complement our reference-based approach and document high within-strain diversity of ADS operon structure and activity in health and disease. For example, we found relatively consistent and high expression of ADS genes among the majority of transcripts assigned to *Leptotrichia* (Fig. 5; Table S11) while expression levels among *Streptococcus sanguinis* and *parasanguinis* is more variable (Fig. 6; Table S12). The results of this analysis demonstrate the diversity of the ADS operon among oral taxa, particularly in understudied groups like *Leptotrichia*.

**FIG 5.**
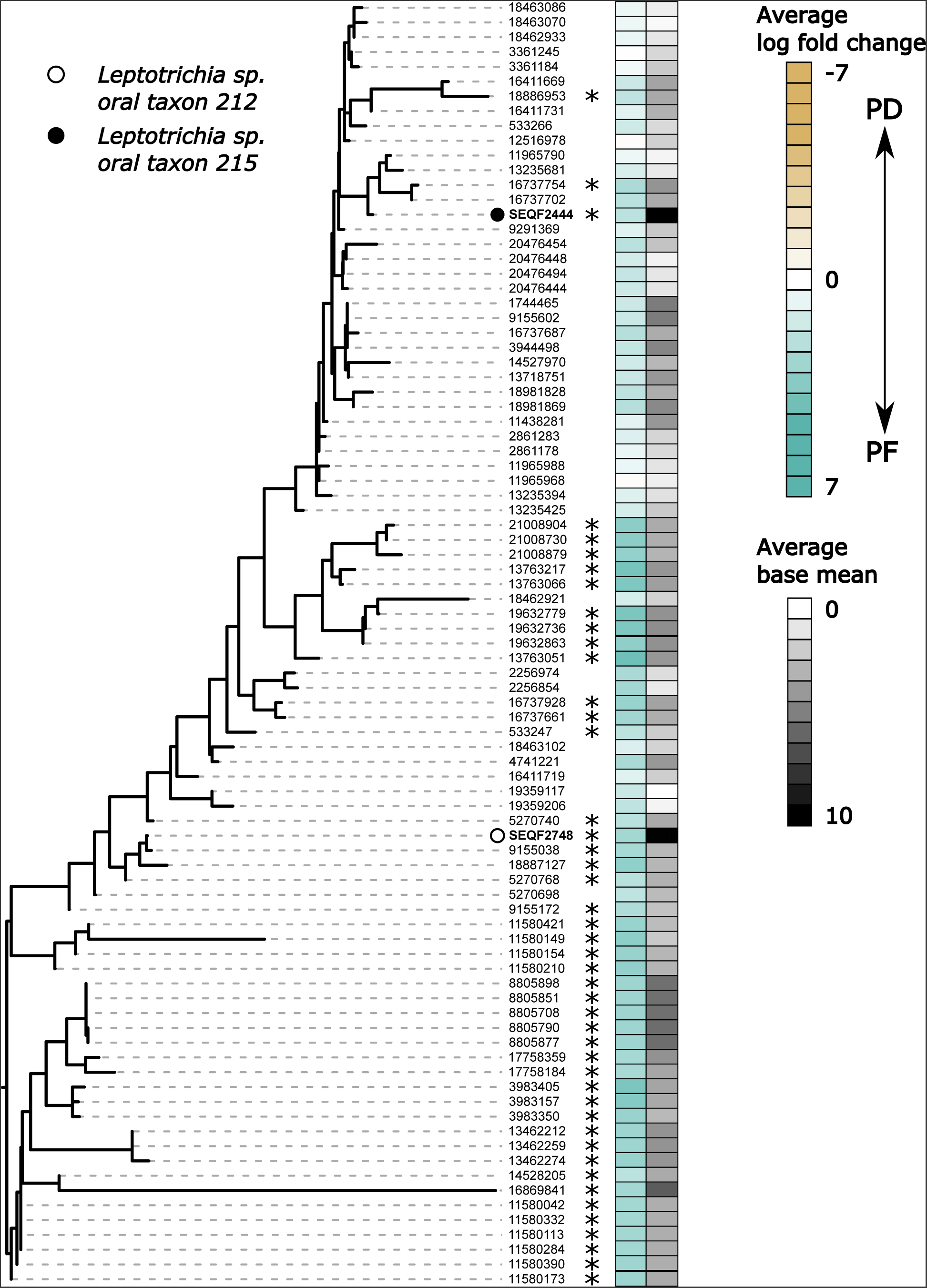
*Denovo* assembly of ADS transcripts document high operon diversity among oral *Leptotrichia*. Maximum likelihood tree of reference HOMD *Leptotrichia* (SEQF, bolded on tree) and *denovo* assembled ADS transcripts assigned to these species (numbers only). Heatmap corresponds to the average log fold change over all three arc genes and average base mean based on either HOMD reference genome mapping (SEQF lineages) or *denovo* assembly (numbered lineages). All lineages have relatively high gene expression in PF with many lineages having all three genes significantly up regulated (denoted by asterisks).

**FIG 6.**
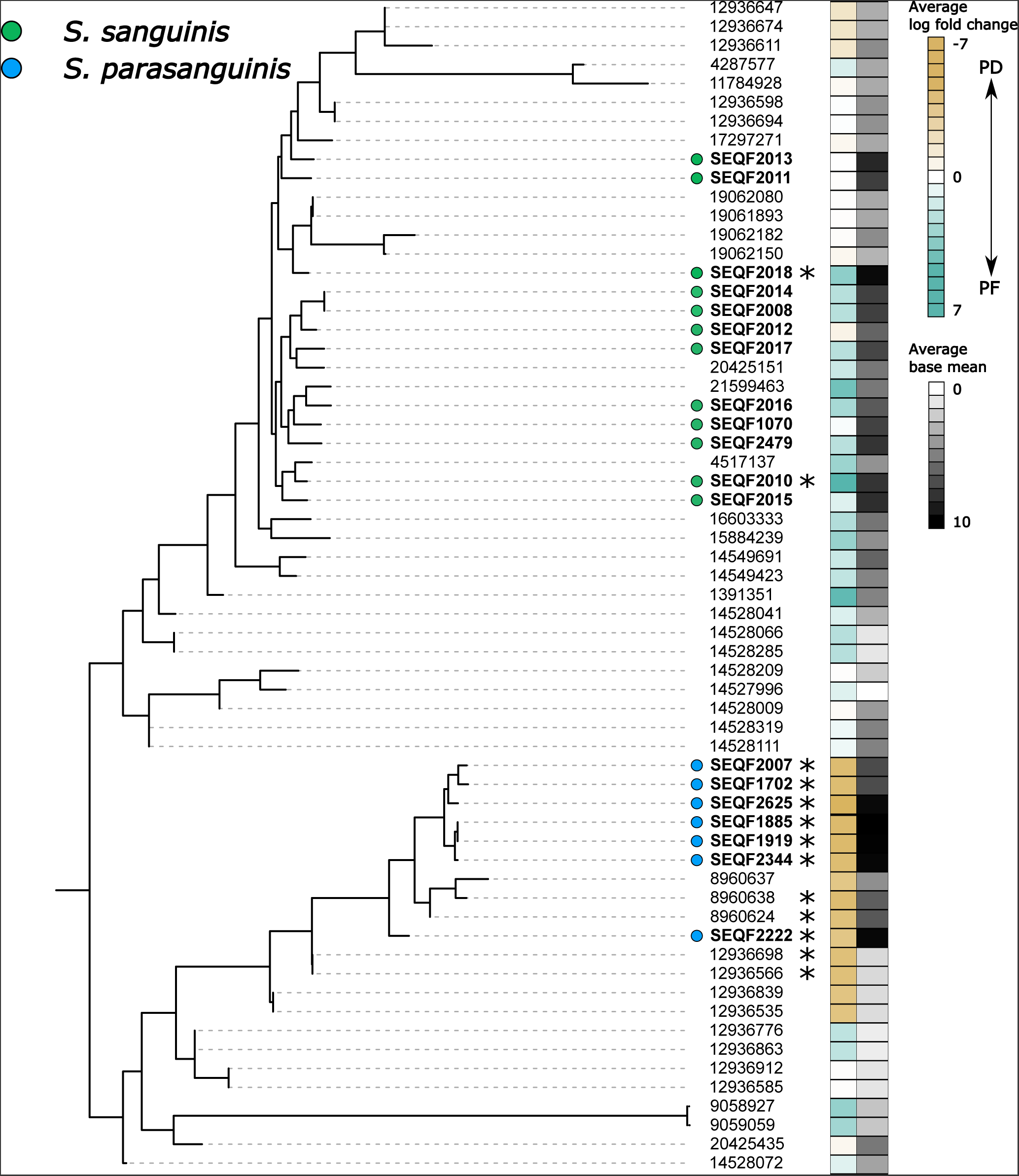
*Denovo* assembly of ADS transcripts document variable expression in health and disease among two Streptococcus species. Maximum likelihood tree of reference HOMD Streptococcus found in a subset of Figure 3 (*S. sanguinis* and *S. parasanguinis*; bolded SEQF annotations) and *denovo* assembled ADS transcripts assigned to these species (numbers only). Heatmap corresponds to the average log fold change over all three arc genes and average base mean based on either HOMD reference genome mapping (SEQF lineages) or *denovo* assembly (numbered lineages). Within species there is variable expression levels among the two species in health and disease with only a subset having all three genes up or downregulated (denoted by asterisks).

### *Streptococcus mutans* antagonism

Next, as ADS activity and ADS competent bacteria have previously been shown to reduce the ability of *Streptococcus mutans* to proliferate within the plaque biofilm, we compared the proportions of ADS competent bacteria and *S. mutans* in both PD and PF samples. We found that while PD samples as a whole had a higher prevalence across samples (77.78%) and overall community proportion (12.43% ± 18.47%) of *S. mutans* as compared to PF (prevalence: 46.88%, average proportion: 2.40% ± 8.44%), 55.56% had low (<5%) *S. mutans* abundance (Fig. 7a) and *S. mutans* abundance was not correlated with citrulline production (Fig. S1). Next, we performed selbal analysis (Rivera-Pinto et al. 2018) to identify groups of taxa that are most associated with citrulline production and found that the balance in ratio between *Veillonella parvula* and *Corynebacterium matruchotii* has the highest association with the total amount of citrulline detected in our samples (Fig. 7b; R^2^=0.32).

**FIG 7.**
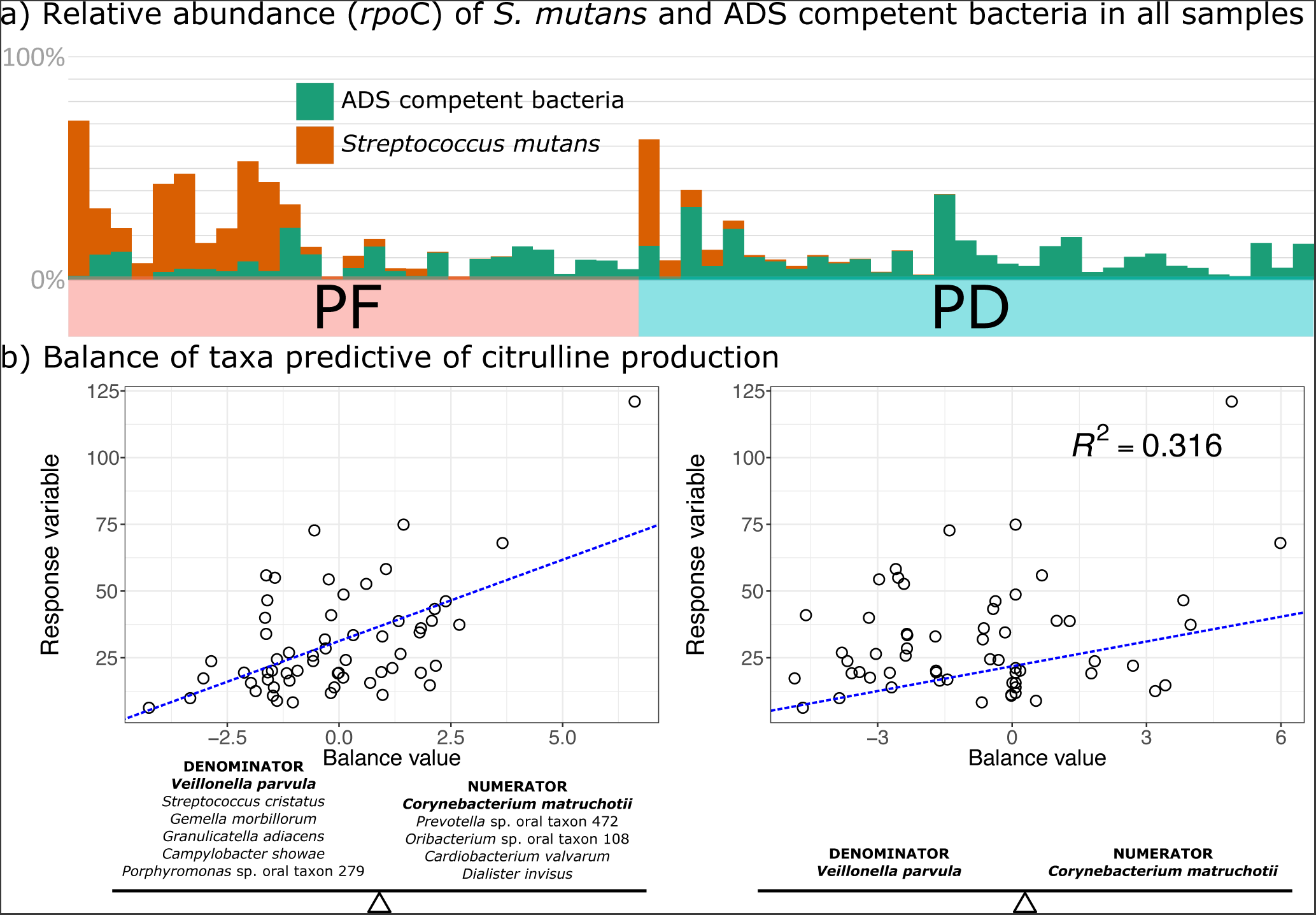
Evidence for Streptococcus mutans antagonism with ADS competent bacteria. (a) Relative proportion of amplicon reads for ADS competent bacteria identified in our metatranscriptomic dataset as compared to *Streptococcus mutans* across all samples sequenced. (b) Balance of taxa most predictive of citrulline production. Scatter plot on the left illustrates the maximum number of taxa whose ratio is associated with the response variable. Plot on the right illustrates the two taxa whose ratio best predicts the maximum amount of variation in the response variable.

Finally, samples with low *S. mutans* (<10% of the total community), have upregulation of the ADS pathway for a variety of taxa including *Parvimonas micra, Actinomyces naeslundii, Bulleidia extructa, Fusobacterium* sp. HMT 370, *Lactobacillus ultunensis, Olsenella uli, Peptostreptococcaceae nodatum, Streptococcus constellatus*, and *Treponema vincentii*. We found no significant correlation between the relative abundance of these taxa and *S. mutans* in our ASV dataset. Upregulation of the ADS pathway is disproportionately skewed by PD samples with low *S. mutans* prevalence and only a single gene assigned to *Streptococcus infantis* (*arc*B) is upregulated in PD samples with high *S. mutans* (Fig. S2a; Table S13). Overall, samples with high *S. mutans* abundance also have a lower relative abundance of *S. oralis* and *S. sanguinis* with the exception of two PF samples (Fig. S3). Interestingly, both PF samples (UF17PF, UF55PF) with a relatively high abundance of *S. mutans* also have a large proportion of total ADS transcripts assigned to *S. sanguinis*. For example, 48% of *rpo*C amplicons derived from sample UF55PF were assigned to *S. mutans* while 14% were assigned to *S. sanguinis*, yet 52% of all ADS transcripts originating from this sample were assigned to *S. sanguinis*.

### Stress response and virulence genes are upregulated in disease

Finally, while this study focused on gene expression for a single metabolic pathway, arginine deiminase, we found 210,295 genes that had significantly different expression between PD and PF samples, 63,268 (30%) of which are unknown hypothetical proteins. Much of the functional diversity at the community level is driven by PD samples which exhibit low functional and community cohesion as compared to the PF samples (Fig. 8a). These differences are the result of a strong functional signal of cariogenic taxa (and particularly *Streptococcus mutans* and *Propionibacterium acidifaciens* (also known as *Acidipropionibacterium acidifaciens,* Fig. 8b). Genes highly expressed in PD samples as compared to PF include a variety of stress response or virulence factors including those involved in biofilm formation, acid tolerance, the production of antimicrobial peptides, and response to environmental stressors. For example, *pfl*B (pyruvate format-lyase-encoding) genes, are key factors in the colonization of *Salmonella* in the gut facilitated by host cell apoptosis (Anderson et al. 2021), *clp*C encodes an ATPase involved in response to environmental stress and proteolytic activity (Biswas et al. 2021; Miethke, Hecker, and Gerth 2006), *dna*K protects against a wide variety of environmental stressors and promotes biofilm growth and lactic acid fermentation at high temperatures in lactic acid bacteria (Abdullah-Al-Mahin et al. 2010; Jayaraman and Burne 1995; Lemos, Luzardo, and Burne 2007), *adh*E promotes the production of mutacin by *S. mutans* that has antagonistic activity against a wide range of gram-positive bacteria including other members of *Streptococcus* (Merritt and Qi 2012; Tsang et al. 2006), and genes encoding fimbriae (Fimbrial subunit type 1) are involved in biofilm development. Interestingly, downregulation of genes encoding the major and minor fimbriae (*fim*A, *mfa*1) have previously been shown to be disrupted by ADS activity in the oral cavity (Cugini et al. 2013).

**FIG 8.**
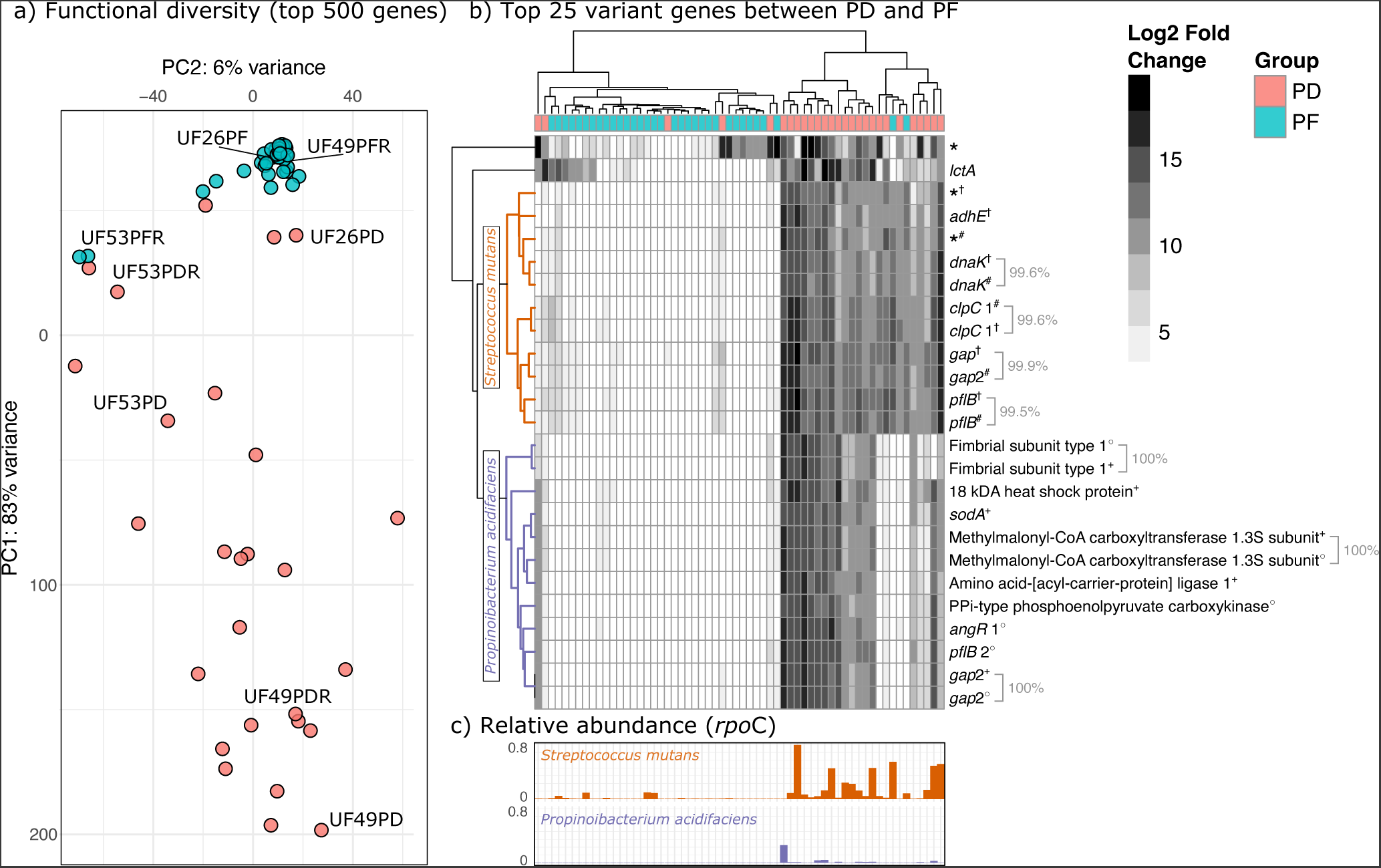
Community-level differential gene expression profiles are more individualistic in PD as compared to PF samples. (a) PCA plot showing the differences in functional profiles among samples using the top 500 varying genes between groups. Teeth sampled from the same individual are highlighted in the plot. (b) Hierarchically clustered heatmap of the top 25 genes (determined by highest variance) that differentiate between PD and PF samples. Genes marked by an asterisk are annotated as hypothetical proteins. Different lineages of *S. mutans* (SEQF1069^†^, SEQF1382^#^) and *P. acidifaciens* (SEQF1851^+^, SEQF2583°) are indicated by corresponding symbols. Identical genes detected in different lineages are indicated by brackets and corresponding percentages indicate genetic identity between the two genes as determined by blastn alignment. (c) Relative abundance of *Streptococcus mutans* and *Propionibacterium acidifaciens* for each sample corresponding to heatmap dendrogram.

We also find multiple *Streptococcus mutans* genes upregulated in disease that are key factors in carbohydrate metabolism including *ccp*A, *cod*Y, *pac*L, *gtf*A, *gtf*C, *lev*D, and *fru*A (Abranches et al. 2008; Moye, Zeng, and Burne 2014). Transcripts assigned to these genes derived from two strains of *S. mutans:* UA159 and NN2025, which are also the dominant *S. mutans* strains found in the *rpo*C dataset (44% and 52% of all *rpo*C amplicons assigned to *S. mutans*, respectively). Like our ADS pathway results, the abundance *of S. mutans* appears to be driving most of the expression level data (Fig. 8c) where 78% of PD samples have *rpo*C reads assigned to *S. mutans* as compared to 48% of PF samples. Interestingly, though, *Propionibacterium acidifaciens* is found at very low abundance across the dataset, even for those samples with high *P. acidifaciens* gene expression. 59.26% of PD and 0% of PF samples had *rpo*C amplicons assigned to this species with an average abundance of 1.25% (± 4.29%) in PD. Similarly, on average 1,088 transcripts (± 1,870) were assigned to *P. acidifaciens* in PD samples as compared to 0.10 (± 0.09) in PF. Assuming this is not the result of the disproportionate representation of *S. mutans* as compared to *P. acidifaciens* in our database (247 versus 2 representative genomes), these results may indicate that this species may be an important driver of tooth decay, even at low frequency in the oral cavity. Importantly, two strains assigned to both *S. mutans* and *P. acidifaciens* are represented in Fig. 8b and contribute relatively equally to the variance between PF and PD as genes found in these strains are too similar or identical to one another to distinguish between genome of origin (ranging from 99.5% to 100% genetic identity).

## DISCUSSION

Manipulation of ADS activity in the oral cavity either through the administration of prebiotics in the form of arginine toothpastes or the development of ADS competent probiotic strains (Acevedo et al. 2005, 2008; Nascimento et al. 2009, 2014; Nascimento 2018; Nascimento, Alvarez, Huang, Browngardt, et al. 2019b; Nascimento et al. 2013; Wolff and Schenkel 2018) are promising prophylactic treatments for the prevention of tooth decay which remains a major global public health problem (Petersen et al. 2005). The results of this study complement and build upon previous investigations of the significance of ADS activity in the oral cavity in preventing tooth decay by providing a multivariate analysis of differential gene expression of ADS competent bacteria in the context of a mixed microbial ecosystem.

In agreement with previous studies (Nascimento et al. 2014; Nascimento, Alvarez, Huang, Hanway, et al. 2019b; Nascimento et al. 2013), we found that while not completely lacking ADS activity, PD samples generally had lower citrulline production from arginine in vitro (Fig. 1b), and a lower proportion of ADS competent bacteria compared to PF samples as measured by both amplicon frequency and transcript frequency (Fig. 5c, 1d). Interestingly, however, we detected thirteen strains of bacteria with upregulation of the ADS operon in caries disease. Most strains identified as upregulated in PD make up a very small proportion of the total oral community (e.g., *Limosilactobacillus fermentum, Lactobacillus ultunensis*) which may indicate that to be effective in preventing tooth decay, the ADS operon must be expressed by a taxon or group of taxa that reach a certain abundance threshold. Alternatively, some taxa may have higher ADS activity in PD but not secrete sufficient levels of ammonia at an adequate rate into the surrounding biofilm matrix to negate environmental acidification, independent of community abundance. Previous research has demonstrated that oral bacteria must be able to generate alkali products above a minimum threshold to prevent acidification of the environment by sugar metabolism (Clancy and Burne 1997) and caries development in experimental animals (Clancy et al. 2000). Not all ammonia produced by alkali-generating metabolic pathways is rapidly released into the extracellular environment. Consider, for example, that while *Streptococcus mutans* lacks the ADS operon, it catabolizes agmatine to produce ammonia, CO_2_, and ATP through the Agmatine Deiminase (AgDS) pathway, which closely resembles arginine catabolism through the ADS. AgDS contributes to acid tolerance and growth of *S. mutans* by ATP and intercellular ammonia production, the latter of which increases cytoplasmic pH, apparently without a substantial effect on the pH of the plaque biofilm (Griswold, Jameson-Lee, and Burne 2006). Moreover, as the ADS pathway is highly regulated, specific environmental conditions may be necessary to trigger higher ADS activity in some species that is not conducive to the prevention of tooth decay. For example, low pH increases the transcription of the ADS pathway in *Streptococcus gordonii* (Liu et al. 2008). It may be the case that the ADS pathway is upregulated at low pH in some taxa where cellular survival is possible, but not the prevention of enamel demineralization. ADS activity can persist in some bacterial species (e.g., *Streptococcus*) at extremely low pH (Marquis et al. 1987) and thus higher transcript expression of the ADS operon may continue even in highly acidic environments. As tooth demineralization is a consequence of not only environmental acidification but the duration of a low pH environment in the plaque biofilm, potential preventative strains of bacteria should have higher ADS expression independent of environmental conditions excluding the availability of arginine itself (Velsko et al. 2018). Finally, our ADS operon coverage analysis and *denovo* assembly of full ADS transcripts suggests that regions of the operon may undergo differential post-translational regulation which may in turn lead to lower ammonia production in some strains or perhaps in certain environmental contexts (Hui, Foley, and Belasco 2014). Further research on the regulatory mechanisms and metabolic behavior of the ADS competent bacteria in PD samples detected here would clarify their efficacy in the prevention of tooth decay.

While we find ADS activity upregulated in both health and disease, it is important to note that upregulation of the ADS pathway in PD samples was limited to those with low (<5%) *Streptococcus mutans* abundance which may suggest that ADS activity prevents the proliferation of this cariogenic taxon in the oral cavity or, alternatively, that the presence of *S. mutans* inhibits ADS activity. Moreover, the oral environment across samples in both health and disease may be more or less conducive to ADS activity. For example, ADS activity is dependent on the availability and concentration of arginine as a secondary carbon source, and can in some taxa be repressed by the oxygen tension in particular sites in the oral cavity (Dong, Chen, and Burne 2004; Zeng, Dong, and Burne 2006). Additionally, coverage plots and *denovo* assembly of ADS transcripts generated in the current study suggest the presence of gene-specific regulation of the ADS operon among certain groups, particularly of the *arc*C gene in *Streptococcus* sp. This post-transcriptional regulation may result from competition for substrates necessary for other metabolic pathways in certain environmental conditions. For example, anaerobic growth of *Streptococcus thermophilus* is strongly dependent on carbamoylphosphate synthetase activity (CPS) which synthesizes carbamoyl phosphate from glutamine, CO_2_, and two ATP molecules as a precursor for arginine and pyrimidine biosynthesis (Arioli et al. 2009). The CPS synthetase domain is found in nearly all organisms (Lawson, Charlebois, and Dillon 1996), and its activity is strongly inhibited by arginine (Arioli et al. 2009; Charlier and Glansdorff 2004; Cunin et al. 1986); thus, in anaerobic environments where arginine is depleted and CPS activity is high, *arc*C activity may be suppressed through selective mRNA degradation to prevent catabolism of carbamoyl phosphate to ATP and NH_3_, instead favoring its use as a substrate for arginine and pyrimidine production. Importantly, in taxa lacking CPS, the arginine deiminase pathway can replace CPS as the primary provider of carbamoyl phosphate for pyrimidine biosynthesis (Nicoloff, Hubert, and Bringel 2001). Finally, higher abundance of ADS competent bacteria may lead to nutritional resource competition with *S. mutans* and other cariogenic bacteria that require arginine and arginine-containing peptides, all of which may contribute to antagonism between *S. mutans* and ADS competent bacteria in both PF and PD. Given that oral health exists on a continuum and tooth decay is a progressive disease, the relative contribution of the oral environment, microbial community structure, environmental secretion of alkaline products of metabolism, post-transcriptional mRNA regulation, and presence or absence of specific strains in promoting health or disease likely fluctuates over time.

Despite this variability, we identified lineages for *Streptococcus oralis, S. sanguinis, S.* sp. HMT 056, and *Leptotrichia* sp. HMT 212 & 215 that may contain effective probiotic strains. Moreover, we detect taxa that may have important roles in the structural development of plaque, the inclusion of which may be necessary for probiotic success. For example, previous research identified *Corynebacterium matruchotii* as one of the taxa that increased with regular brushing with fluoride+arginine toothpastes while species of *Veillonella* decreased (Carda-Diéguez, Moazzez, and Mira 2022). In the current research, we find both *C. matruchotii* and *Veillonella parvula* to be associated with high or low citrulline production, respectively. In addition, *C. matruchotii* plays a significant role in biofilm formation, can induce the mineralization of plaque, and recently was shown to increase the growth rate of various *Streptococcus* sp. *in vitro* (Almeida 2022; Q. Li et al. 2022; Morillo-Lopez et al. 2022). It therefore may be a key structural taxon for the colonization and proliferation of ADS competent bacteria in the oral cavity.

Importantly, the analysis of multiple *Streptococcus* genomes for each species highlighted that ADS upregulation is heterogeneous among lineages identified by our metatranscriptomics dataset (e.g., *S. oralis* = 25%, *S. sanguinis* = 17%), that not all lineages annotated by HOMD possess the full ADS operon (e.g., *S. oralis =* 25%, *S. sanguinis =* 92%. Table S3), and that *denovo* assembly of the ADS operon can identify multiple distinct transcripts for the ADS operon assigned to a single species. Additionally, we found that relatively high proportions of ADS competent bacteria do not equate lower proportions of cariogenic bacteria in all cases. For example, two PF samples in our dataset had both high frequency of *S. mutans* and *S. sanguinis* as measured by *rpo*C sequencing (Fig. S3). However, as these samples also had high ADS activity by *S. sanguinis* one possible interpretation is that this activity has prevented the development of a cavity in these teeth, despite the presence of *S. mutans.* Alternatively, samples with high relative frequency of *S. sanguinis* and *S. mutans* but low ADS activity for *S. sanguinis* (e.g., UF68PD) may reflect strain level differences in ADS competency. Given that some lineages of *S. sanguinis* do not have a full ADS operon (Table S3), it may be that some strains are not capable of antagonizing *S. mutans* even at relatively high frequency. The importance of strain level resolution in determining candidate probiotics to promote ADS activity in the oral cavity has previously been described among strains of closely related *Streptococcus* species (Velsko et al. 2018). Our results confirm the importance of strain level upregulation of ADS in the context of oral health for different *Streptococcus* species (e.g., *S. oralis, S. sanguinis, S.* sp. HMT 056) and identifies potential candidate species that are as of yet understudied for their role in the promotion of oral health. For example, we found two strains (lineages) of *Leptotrichia* (HMT 212 & HMT 215) that had high ADS activity in PF and comparatively high read abundance and prevalence across samples in the current dataset. Moreover, *denovo* assembly of full ADS operons documents high levels of diversity within this group that may contribute to health or disease. Conversely, our results support the suggestion that lineages of bacteria that are typically associated with health (e.g., non-mutans *Streptococcus* sp.) may be important contributors to the etiological development of caries (van Ruyven et al. 2000; Sansone et al. 1993). Along with potential probiotic uses, identification of specific bacterial lineages consistently associated with health and with known beneficial properties in the oral cavity may be used as additional markers of risk assessment beyond the presence of cariogenic taxa, though further investigation into the prevalence and functional activity of the taxa identified here is necessary to clarify their role in maintaining favorable pH in the oral cavity.

Analysis and interpretation of metatranscriptomic datasets generated from mixed microbial communities is complicated by differences in community composition and membership (Baker et al. 2021; Gross et al. 2012), particularly when the microbial community found in comparative groups are highly dissimilar. Thus, accounting for differences in community composition between groups is critical (Klingenberg and Meinicke 2017; Zhang et al. 2021). Here we compared log-fold changes in a parallel metataxonomic dataset to investigate the impact of community composition and species abundance on our metatranscriptomics results. Our paired metatranscriptomic and metataxonomic results demonstrate that often higher gene expression is a consequence of differential community membership or abundance and not truly neutral differences in gene expression profiles. For example, all three *Streptococcus* species found here to have higher ADS activity on healthy teeth typically make up a much larger proportion of PF communities and so more transcripts may be a simple function of more cells contributing to the effect. Conversely, taxa found at low frequency may be functionally more active in some circumstances (e.g., *Limosilactobacillus fermentum, Lactobacillus ultunensis, Propionibacterium acidifaciens*). Given that a good candidate probiotic will need not only to efficiently catabolize arginine to produce ammonia but also have the ability to effectively colonize the plaque biofilm, comparing genome and transcript frequency is an important consideration. Moreover, transcript abundance alone does not necessarily precisely correlate to actual *in vivo* biochemical activity and a variety of confounding factors including strain level translational efficiency and enzyme stability may impede our accurate interpretation of activity levels alone. For the purposes of this study, we focus on taxa with both high RNA transcript abundance and comparatively high ASV abundance as these taxa have (1) high ADS expression that is not directly linked to high relative abundance in a sample and (2) are prevalent across multiple individuals (and thus may be more efficient colonizers of the plaque biofilm).

Finally, we find that while Amplicon Sequence Variant (ASV) diversity is lower in PD as compared to PF samples (Fig. 1a), functional diversity is much more dispersed as compared to healthy teeth which tend to have a much more consistent functional profile (Fig. 6a). The main functional differences between PD and PF communities are largely driven by a variety of genes associated with virulence factors and carbohydrate metabolism in known cariogenic taxa and these effects are largely individualistic in scope which suggests the presence of high functional volatility during caries development. Interestingly, we detected two major taxa driving the majority of differentially abundant genes in PD versus PF, *Streptococcus mutans* and *Propionibacterium acidifaciens,* even though *P. acidifaciens* occurs at low frequency for both groups in our corresponding amplicon dataset. While this may be the result of taxonomic biases inherent in amplicon datasets, it may also demonstrate important functional activity by relatively low frequency taxa. Moreover, as *P. acidifaciens* biofilm formation is inhibited by *S. mutans* growth (Obata et al. 2019), a better understanding of the functional interaction between these species may elucidate their impact on the total ecology. The volatility of the functional profile of PD teeth coupled with lowered diversity suggest that candidate probiotics should not be limited to a single species.

Results of the current study highlight potential candidates for probiotic panels including oral bacteria where pH modulation through the ADS pathway have previously been identified (e.g., *Streptococcus* sp.) (Burne and Marquis 2000) as well as those that are less well characterized (e.g., *Leptotrichia* sp.). In conclusion, while our multivariate approach substantiates the role of the ADS pathway in health and disease, it highlights the importance of accounting for taxonomic shifts in the analysis of functional differences in mixed microbial ecosystems such as dental plaque, which will impact the efficacy of any caries treatment and suggests that probiotic development may benefit from further strain-level investigation of a wider assortment of species for ADS activity and pH modulation potential in mixed microbial communities.

## MATERIALS AND METHODS

### Samples and study design

A total of 54 adult individuals were recruited for this study. Carious lesions were diagnosed using the International Caries Detection and Assessment System II (ICDAS-II) visual criteria (Ekstrand et al. 2007) and the lesion activity (LAA) (Braga et al. 2009) scoring system by a calibrated examiner (M.M.N.) as previously described (Nascimento et al. 2013; Richards et al. 2017). Sampling of site-specific plaque samples followed protocols previously published by this group (Nascimento et al. 2013b). Briefly, study participants were instructed to refrain from oral hygiene procedures for 8 hours prior to plaque sampling, after which supragingival plaques were collected from individual teeth and classified as caries-free (PF) (ICDAS score = 0) or with an active dentin carious lesion (PD) (ICDAS score = ≥4). PD samples were recovered from the internal surface of the lesion. Full metadata for each sample can be found in Table S14.

### Citrulline assay

A total of 74 supragingival plaque samples were tested for their potential to generate citrulline from arginine via the arginine deiminase system using a protocol previously validated and published by our group (Velsko et al. 2018). Briefly, plaque samples were dispersed using external sonication for two cycles of 30 seconds with cooling on ice during the interval. The cells were then harvested, washed, and resuspended in 10 mM Tris-Maleate buffer and further permeabilized with toluene-acetone (1:9) for the measurement of ADS activity. The total protein concentration of the plaque sample was also measured using a BCA protein estimation kit (Pierce, Waltham, MA, USA) with a known concentration of bovine serum albumin (BSA) as a standard. The ADS activity levels in the plaques were normalized to protein content and represented as nanomoles of citrulline generated per milligram of protein.

### DNA and RNA extraction

Plaque samples were stored in RNAlater^TM^ (ThermoFisher Scientific, Invitrogen, Waltham, MA) immediately after sampling. DNA and RNA were extracted simultaneously following the mirVana miRNA isolation kit for RNA protocol (Ambion, ThermoFisher Scientific, Waltham, MA). We chose this isolation kit as it is designed to efficiently isolate small RNAs (<200 nt) as well as total RNA which play a crucial role in regulating bacterial global gene expression and is routinely used for microbiome studies (Azpiroz et al. 2021; Tong, Tang, and Wang 2023; Yost et al. 2018). For RNA isolation, genomic DNA was removed using a Turbo DNA-free kit (Applied Biosystems, Ambion). For DNA isolation, the organic phase from the mirVana mRNA isolation was stored at 4°C until use. An extraction buffer containing 0.1 M NaCl, 10 mM Tris-HCL (pH 8.0 ± 0.2), 1 mM EDTA, and 1% SDS was prepared. The final pH of the buffer was adjusted to pH 12 (± 0.2) with 10 N NaOH. We next added 250 µl of the extraction buffer to the organic phase containing DNA followed by an incubation at 4°C. After centrifugation at 10,000 g for 20 minutes, the aqueous phase was recovered and immediately stored at −80°C for an hour after addition of 1/15 volume of 7.5 M ammonium acetate and 100% ethanol. The samples were then centrifuged for 20 minutes at 10,000 g and the supernatant was discarded. The pellets were air dried and washed with ethanol before being resuspended in 17 µl of molecular grade water.

### RNA library preparation and sequencing

Total RNA samples were processed twice with MICROBEnrich^TM^ and MICROBExpress^TM^ bacterial mRNA enrichment kits (Ambion/Life Technologies, Grand Island, NY) to remove eukaryotic mRNA and 16S/23S rRNAs, respectively. The purified mRNA was resuspended in 25 µl of nuclease-free water. The quality of enriched mRNA was analyzed using an Agilent Bioanalyzer (Agilent Technologies, Santa Clara, CA). Next, we generated cDNA libraries from 100 ng of enriched mRNA using the NEBNext Ultra directional RNA library preparation kit and NEBNext dual index oligonucleotides for Illumina (New England BioLabs, Ipswich, MA). To ensure proper enrichment of bacterial transcripts we first sequenced the pooled library using a MiSeq Reagent Nano Kit v2 (300-cycles) at Clemson University. Deep sequencing of the library was next performed by the NextGen DNA sequencing core laboratory of the Interdisciplinary Center for Biotechnology Research at the University of Florida (Gainesville, FL). Samples were sequenced on the NovaSeq 6000 platform on a single S4 flow cell using a paired end 2×150 bp chemistry.

### DNA library preparation, and sequencing

Each plaque sample was amplified and built into Illumina sequencing libraries using a custom primer set targeting an approximately 478 bp fragment of the bacterial *rpo*C gene - rpoCF (5’-MAYGARAARMGNATGYTNCARGA-3’) and rpoCR (5’-GMCATYTGRTCNCCRTCRAA-3’) (Mann et al. 2023). We adopted this approach as recent research has demonstrated the inability of other common metabarcoding approaches (e.g., 16S rRNA hypervariable regions) for resolving common oral groups (i.e., *Streptococcus* sp.) (O’Connell et al. 2022) and further research identified the *rpo*C gene as a promising alternative marker (Hassler et al. 2022). We note, however, that like other amplicon-based approaches, *rpo*C gene fragment metabarcoding is not free from bias (see (Mann et al. 2023)) and the primers published here have only been tested using oral samples. Each 25 µl PCR reaction consisted of 0.5 µl of the forward and reverse adapter, 10 µl of H_2_0, and 10 µl of Platinum Hot Start PCR Master Mix (2X) (Invitrogen, Carlsbad, CA). PCR thermocycler conditions were as follows: 94°C for three minutes followed by 41 cycles of 94°C for 45 seconds, 39.5°C for one minute, and 72°C for one minute and thirty seconds. A final elongation step was performed at 72°C for 10 minutes. Amplification of PCR products for each sample was confirmed by both gel electrophoresis and Qubit Fluorometer (Invitrogen, Carlsbad, CA). We processed PCR blanks (molecular grade water) in parallel to all samples. Prepared libraries were sequenced on an Illumina MiSeq using 2×300 paired-end chemistry.

### Metatranscriptomic data processing

On average, we generated 326,151,580 (SD ± 102,407,864) raw sequencing reads per sample. We first assessed the quality of the raw reads using FastQC v0.11.9 (Andrews 2019) before quality filtering, trimming adapters, and removing poly-G tails using Cutadapt v3.3 (Martin 2011). On average, 302,238,342 (SD ± 9,238,698) reads passed our quality filtering thresholds. Next, we performed a preliminary filter on the trimmed reads by first mapping to the SILVA v138 non-redundant 99% SSU and LSU 18S rRNA and 16S rRNA databases (Quast et al. 2013) and then to the human reference genome (GRCh38) using Bowtie2 v2.4.5 (Langmead and Salzberg 2012). On average 0.02% (SD ± 0.06%) of our quality filtered reads per sample mapped to the SILVA database and 3.04% (SD ± 4.43%) to the human genome. We removed any reads that mapped to either the human reference genome or SILVA ribosomal databases using a custom Python script before performing downstream analyses.

For taxonomic and functional assignment, all filtered reads were mapped against the Human Oral Microbiome Database (HOMD v9.15) (Chen et al. 2010) using STAR v2.7.5c (Dobin et al. 2013) with alignIntroMax set to one to disable spliced alignments and outFilterMismatchNoverLmax set to 0.05 to increase mapping specificity. On average 198,385,752 (SD ± 68,279,777) reads mapped to our database per sample. Gene counts were calculated from each sample using the featureCounts function as implemented in the Subread v2.0.1 package (Liao, Smyth, and Shi 2013, 2014). Given that our reference-based approach included a database of diverse microbial genomes with varying levels of genomic congruence, when running featureCounts we allowed for multimapping genes (-M), only pairs with both reads aligned were counted (-B) and paired reads that mapped to separate chromosomes (or contigs in the case of incomplete genome reconstructions) were not counted for downstream analyses (-C).

To complement our reference-based approach, we performed an additional *denovo* assembly on a sample-by-sample basis using Trinity (v2.15.1) (Grabherr et al. 2011). On average we generated 364,811 transcripts per sample (± 159,936). We next merged the assembled transcripts and performed a full length dereplication using vsearch (v2.14.1) (Rognes et al. 2016). We annotated the final dereplicated database using prokka (v1.14.5) (Seemann 2014). Finally, we mapped our filtered reads to the *denovo* reference using STAR (v2.7.5c) (Dobin et al. 2013) in an identical manner as described above. We identified significantly up or down regulated genes using DESeq2 (Love, Huber, and Anders 2014). We built maximum likelihood trees of our *denovo* ADS transcripts with references from HOMD using mafft (v7.453) (Katoh et al. 2002) and RAxML (v8.2.12) (Stamatakis 2014), which were visualized in iTOL (v5) (Letunic and Bork 2021). As many transcripts were truncated or otherwise incomplete, we built phylogenetic trees with only those transcripts that had annotations for all three arc genes (arcABC) and trimmed the final alignments using trimAl (v1.4.rev15) with a gap threshold of 0.5, minimum position overlap in a column of 0.5, and minimum percentage of “good positions” a sequence must retain for inclusion of 50 (Capella-Gutiérrez, Silla-Martínez, and Gabaldón 2009) Assembly statistics can be found in Table S8.

### Metataxonomic data processing

On average, we generated 18,417 raw amplicon sequences per sample (± 17,284) of which 11,517 (± 10,347) passed our quality filtering thresholds. Rarefaction curves for all samples can be found in Fig. S4. Briefly, we trimmed primer and adapter sequences from our raw reads using Cutadapt (v4.0) (Martin 2011), after which we quality filtered, trimmed, merged, removed chimeric sequences, and generated ASVs using DADA2 (v1.22.0) (Callahan et al. 2016). In total, we generated 3,928 unique ASVs. We next assigned a taxonomy to each ASV using Kraken2 (v2.1.2) (Wood, Lu, and Langmead 2019) with the Human Oral Microbiome Database whole annotated genomes as reference (Chen et al. 2010). Low frequency ASVs (prevalence threshold = 0.01) or those with no or kingdom-only taxonomic assignments (i.e., Unknown or Bacteria) were removed from downstream analyses leaving 3,114 ASVs in the dataset. A full ASV table with annotated taxonomy can be found in Table S15. All amplicon sequence analyses were performed in the R v4.1.2 environment (R Core Team 2017). We calculated alpha and beta diversity metrics using the Phyloseq (v1.38.0) (McMurdie and Holmes 2013), vegan (v2.6-2) (Oksanen et al. 2019), and ggdist (v3.2.0) (Kay 2022) R libraries. We generated capscale plots using Bray-Curtis dissimilarity and a distance-based redundancy analysis approach (Legendre and Anderson 1999) to visualize differences in beta diversity across samples. Ordination of samples was constrained by tooth type (PD vs PF) and citrulline expression (log2 transformed) (Fig. 1b). Significance between groups was calculated on isometric log-ratio transformed data normalized using PhILR (v1.20.1) (Silverman et al. 2017). Finally, we identified groups of taxa associated with citrulline production using selbal (v0.1.0) (Rivera-Pinto et al. 2018).

### ADS operon identification

The ADS operon was identified in the HOMD database by first selecting all genes annotated as those that produced one of the three main enzymes involved in the pathway. While gene content varies within and among groups the enzymes encoded by three genes: (1) *arc*A (arginine deiminase EC: 3.5.3.6), (2) *arc*B (ornithine carbamoyltransferase EC: 2.1.3.3), and (3) *arc*C (carbamate kinase EC: 2.7.2.2) are necessary for ADS activity and are thus the focus of this manuscript (Griswold et al. 2004; Liu et al. 2008; Noens and Lolkema 2016b; Novák et al. 2016b). Because gene annotations are rarely consistent across published datasets, we first performed homologous gene clustering using the HOMD database to verify arc gene identity using BLAST (Altschul et al. 1990) and the Markov Clustering (MCL) algorithm implemented in MCLblastline (Enright, Van Dongen, and Ouzounis 2002). In brief, we created a BLAST database of all protein sequences that had been annotated as one of the three ADS enzymes (i.e., by the EC number) from the HOMD database. We next mapped the full HOMD protein database against our reference ADS database using BLASTp (evalue 1e-10), the output of which we used to generate homologous gene clusters using MCLblastline with an inflation parameter of 1.8 which previously had been identified as well suited for high-throughput data (Brohée and van Helden 2006) and in practice can identify homologous gene clusters across diverse groups (Townsend et al. 2021). Genes that did not cluster with any other *arc* gene in the database (e.g., singleton clusters) and annotated genes that fell into clusters that included a mixture of genes annotated as *arc*A, *arc*B, and *arc*C were removed from downstream analyses. Using results from this analysis, we further identified species with a full ADS operon as those wherein all three genes (arcA, arcB, and arcC) were (1) present and (2) in synteny with one another in the reference genome. Synteny was manually identified using locus tag assignment. Species where synteny of the arc genes was interrupted by a single gene (e.g., hypothetical protein) were included in downstream analyses to account for operon variability. A full list of species that carry the ADS operon from this analysis can be found in Table S2.

### Differential gene expression and species abundance analyses

We calculated differential gene expression using DESeq2 v1.34.0 (Love, Huber, and Anders 2014) in R v4.1.2 (R Core Team 2017). To minimize the effect of large fold changes between groups that were the result of low transcript count we next ran lfcShrink on our DESeq2 results using the apeglm R package (Zhu, Ibrahim, and Love 2019). Our threshold for differential expression between groups was an adjusted p-value (FDR) of less than or equal to 0.05 and a minimum fold change of four. Because we expect community composition and species presence to be meaningful confounding variables in comparing estimates of gene expression across groups, we calculated the correlation between log fold change values in transcript abundance to *rpo*C gene abundance for a selection of ADS competent oral taxa (Fig. 5c, Table S6).

### Streptococcus ADS operon phylogeny

First, we concatenated the HOMD annotated *arc*A, *arc*B, and *arc*C genes for each *Streptococcus* lineage in the HOMD database that were validated by our homologous gene clustering into a single contiguous sequence using the concat function in SeqKit (v2.3.1). Concatenated sequences were then aligned using mafft (v7.505) (Katoh et al. 2002) and built into a maximum likelihood phylogeny using RAxML (v8.2.12) (Stamatakis 2014). The phylogeny was midpoint rooted and visualized in FigTree (v1.4.4) (Rambaut 2018). Collapsed log fold change and base mean was calculated by taking the average across all three *arc* genes for each *Streptococcus* lineage. Lineages that fell outside of an expected major species clade were checked using the BLAST web application (Altschul et al. 1990) and the full NCBI NT database as reference to ensure that it was annotated with the correct taxonomic assignment.

### ADS operon coverage plots

Using sequence coordinates listed for each *arc* gene in the corresponding HOMD GFF files, we used the BEDTools (v2.27.1) (Quinlan and Hall 2010) function getfasta to pull the *arc*ABC operon for each species of interest. Next, we mapped the full quality filtered metatranscriptomic sequencing data for each sample to the selected operon sequences using Bowtie2 (v2.4.5) (Langmead and Salzberg 2012). Depth of coverage was calculated using SAMtools (v1.12) depth (H. Li et al. 2009) and plotted in R (v4.2.0) (R Core Team 2017).

## DATA AVAILABILITY

Conda environments and processing scripts are provided at https://github.com/aemann01/ads_plaque and archived under the DOI: 10.5281/zenodo.7760990. Raw metatranscriptomic data has been deposited in the European Nucleotide Archive under the accession number PRJEB60355 and *rpo*C amplicon data under accession number PRJEB60856.

## ACKNOWLEDGEMENTS

This research was supported by the National Institutes of Health National Institute of Dental and Craniofacial Research (R01DE25832 to R.A.B.) and in part, by computational resources provided by the Clemson University Genomics and Bioinformatics Facility, which receives support from two Institutional Development Awards (IDeA) from the National Institute of General Medical Sciences of the National Institutes of Health under grant numbers P20GM146584 and P20GM139767. We thank Alejandro Riveros Walker for comments on an early version of this manuscript and discussions on methodology.

