## Supplementary Figures for "Heterogeneous lineage-specific arginine deiminase expression within dental microbiome species"

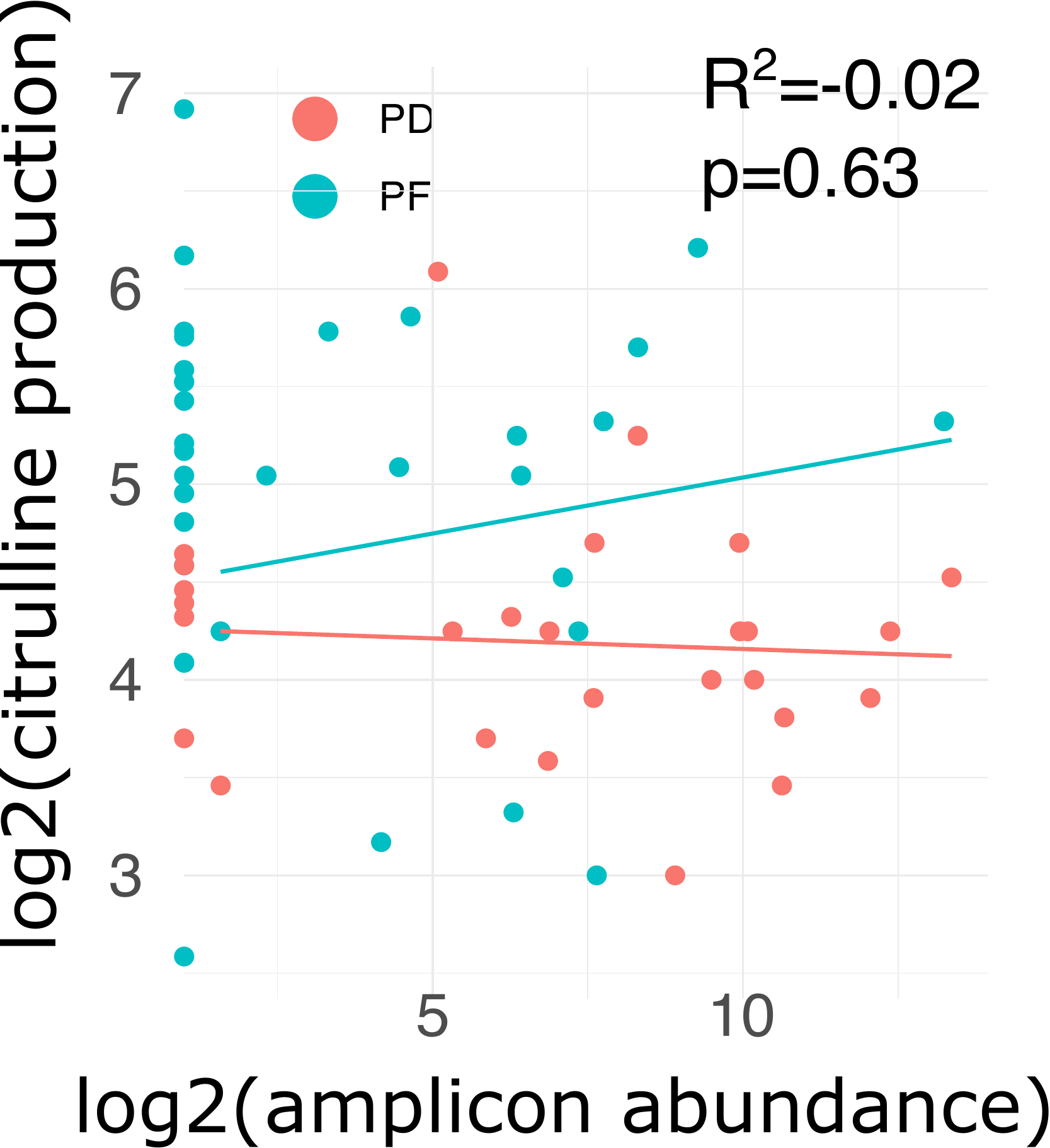

**FIG S1.** Correlation of citrulline production with the abundance of *Streptococcus mutans* as measured by ASV relative abundance. Trend lines are shown for each group (PD or PF), R^2^ value and p-value on graph adjusted for both groups.

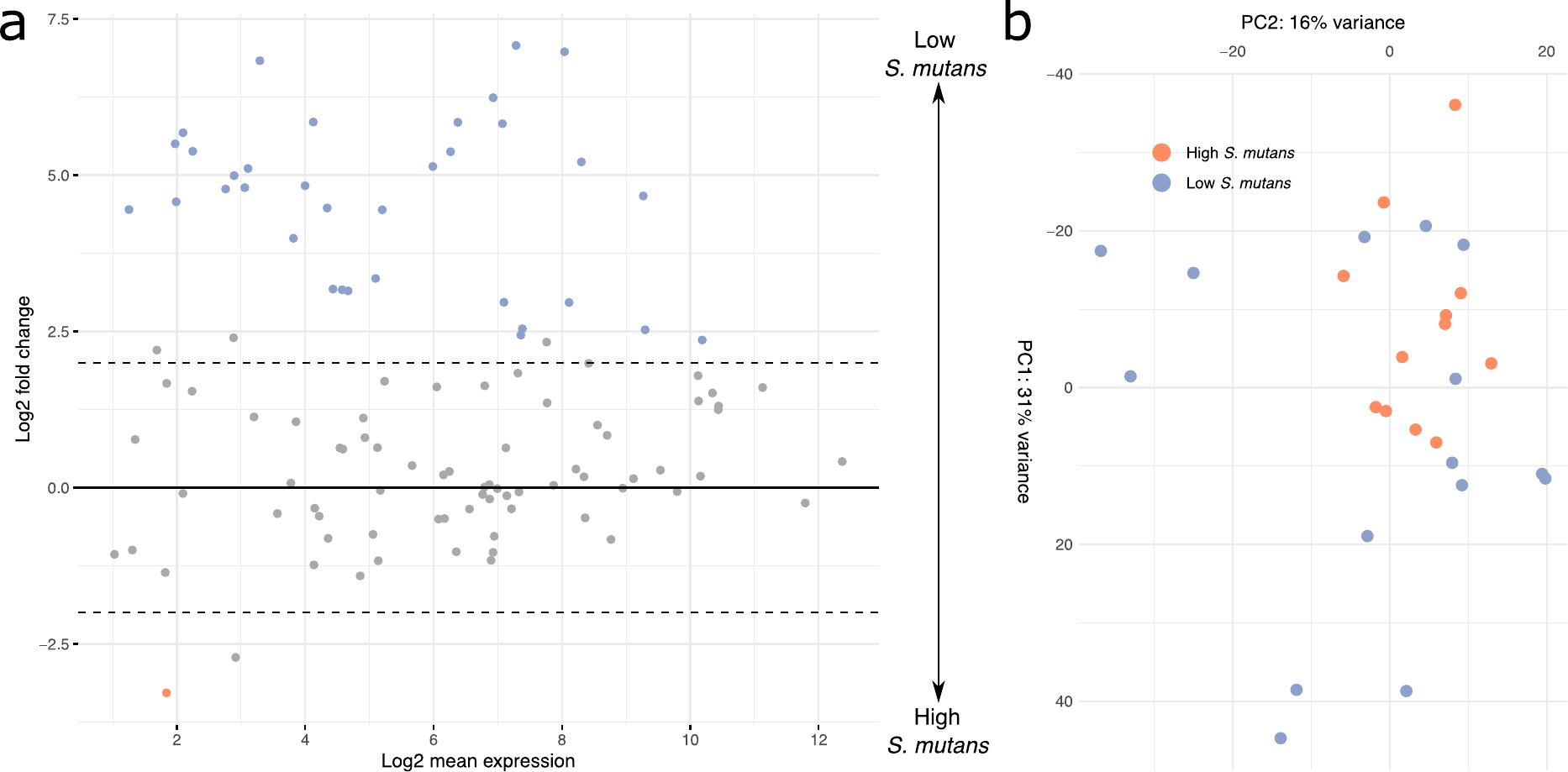

**FIG S2.** (a) Volcano plot of ADS genes (collapsed by species) up or down regulated in PD samples with high (>5%) or low (<5%) abundance of *Streptococcus mutans* as measured by *rpo*C sequencing. (b) Corresponding principal coordinate plot illustrating the ADS gene expression similarity between individual PD samples colored by *S. mutans* abundance group.

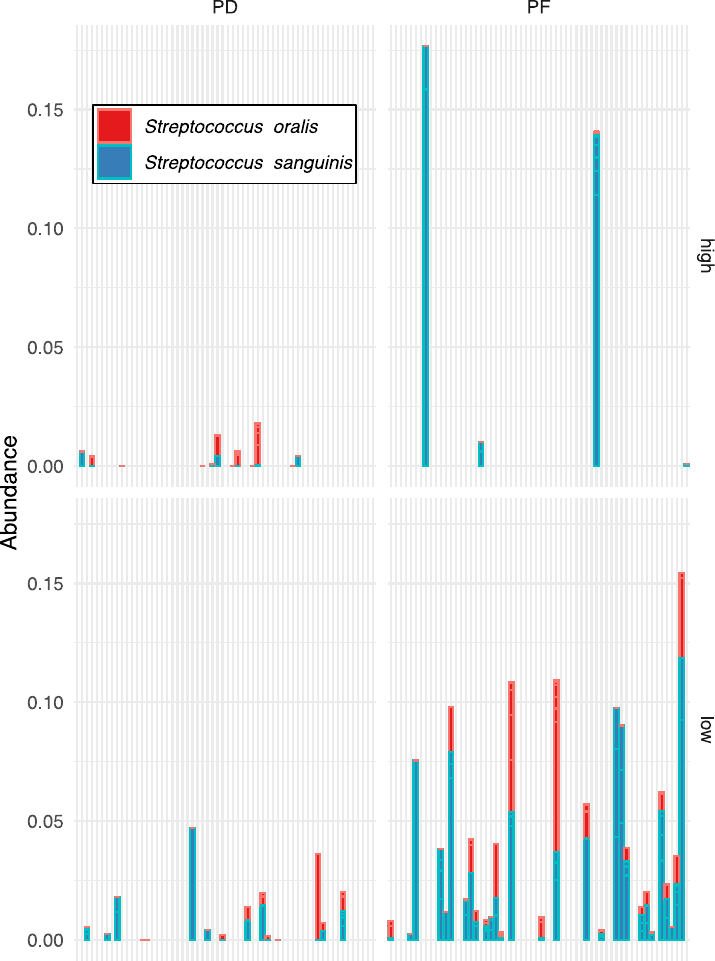

**FIG S3**. Relative abundance of two ADS competent species, *S. oralis* and *S. sanguinis* in PF and PD samples with high (>5%) or low (<5%) *S. mutans* abundance.

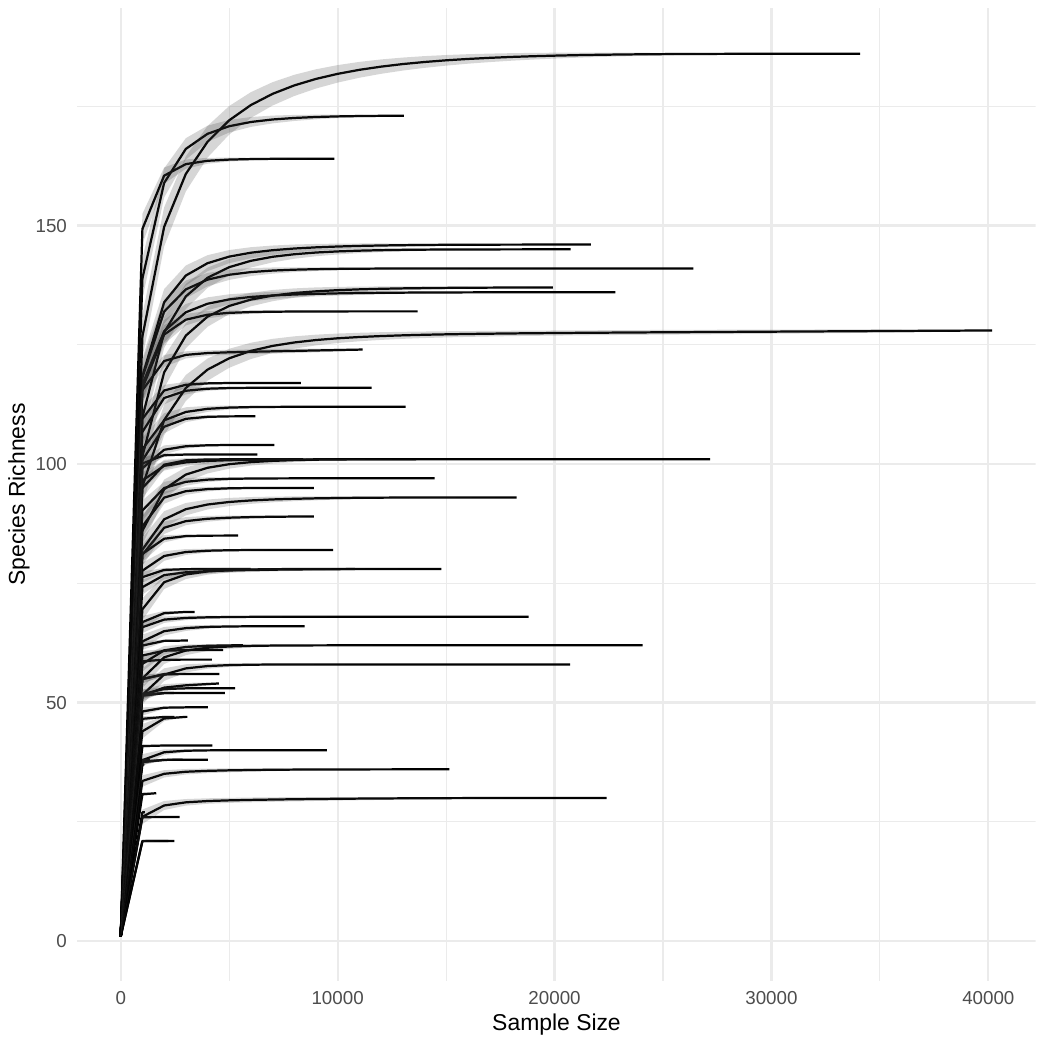

**FIG S4.** Rarefaction analysis of all samples.
